# Autonomous Remotely Controlled Closed System Transgenic Cell Technologies Robot: CRISPR.BOT

**DOI:** 10.1101/2022.07.22.497959

**Authors:** Fatmanur Erkek, Gamze Gulden, Berranur Sert, Menekse Cagla Yilmaz, Sibel Pinar Odabas, Enes Bal, Gamze Yelgen, Tarik Teymur, Yasin Ay, Solen Dogdu, Nulifer Neslihan Tiryaki, Buse Baran, Beste Gelsin, Hasret Araz, Ilayda Cavrar, Cihan Tastan

## Abstract

In manually advancing experimental processes, the stages may be long-term and need to be repeated. Human errors with the repetition of the steps turn into a time-consuming and high-cost for the experiment processes. For this reason, autonomous liquid processing systems are promising technologies. However, in addition to the high cost of fully automatic systems, their maintenance is also quite expensive. Furthermore, conventional systems usually require system-specific protocols and laboratory equipment. Here, we aimed to show that the autonomous robotic systems may provide a closed and error-free molecular biology bench to perform genetic engineering automatically, quickly, and practically 7-24. In this way, researchers can save time from repetitive experiment processes and perform BSL3 experiments including pathogens without human contact. In this study, we built CRISPR.BOT robotic systems to perform Green Fluorescent Protein (GFP) encoding plasmid DNA transfer into bacteria, lentiviral transduction of the gene-of-interests including GFP encoding gene and CRISPR-Cas9 with gRNAs genetic editing system to a human cell line. Furthermore, we showed that CRISPR.BOT system achieved to accomplish single-cell subcloning of GFP+ CRISPR-gRNA+ cells with 90-100% purity. This study suggests that CRISPR.BOT-like approaches may reduce manpower in a safely closed bench in which molecular biology and genetic engineering can be done by robots in a closed system without touching pathogenic microorganisms (virus or bacteria, for example, SARS CoV-2 virus). Furthermore, LEGO Mindstorms robots showed to have the potential to be used in daily laboratory routines with their cost-effectiveness reduced by up to 50 times compared to normal commercial robots.

## Introduction

Liquid-processing systems technology is involved in various fields of life sciences. Automated liquid handling systems are used day by day to develop protocols and for reliability in laboratory fields such as synthetic biology, microbiology, or genetics. LEGO Mindstorms, which is widely used in the programming of complex systems and offers a wide range of work, provides opportunities such as designing machine prototypes in the field of robotics (Guidi, 2015). It has been reported in the literature that liquid-processing LEGO robotic systems can be used in biotechnology experiments (Gerber, 2017). In the future, these studies are thought to be useful for complex drug delivery experiments and many other applications in emerging fields such as regenerative medicine and tissue engineering, and biotechnological applications (Wagner, 2019). In addition to the high costs of fully automatic systems, they are also quite expensive to maintain (Turbak, 2002). It is beneficial to have systems that are affordable and easy to install. However, when we look at the market, there are no reasonable robotic systems that meet the requirements, precise and programmable at the same time. It is beneficial that systems with easy installation and affordable prices are available. Also, in the market, there are no specialized robotic systems that can perform molecular biology and genetic editing techniques in an affordable and precise way. With this project, machine learning, transgenic cell screening, analytical calculations, gene designs, genetic pathway selection, experimental design, cell cloning, and genetic pathway analyzes may be performed with a very low margin of error, without human intervention, by developing remote and autonomous robotic systems.

LEGO Mindstorms robots offer a cost-effective system that can provide the same experimental efficiency as autonomous robotic systems, as well as a programmable and remote-control system. It has been demonstrated that liquid-processing robotic systems like the CRISPR.BOT has the potential to be used in biotechnology experiments and laboratory routines. We have built the prototype of an innovative technology that is cost-effective, practical, and free from human error in bioindustry applications. In our first experimental study, in which we showed all experimental stages of bacterial DNA transformation, production of transgenic human cells with lentiviruses, and CRISPR gene-engineered cell development with developed algorithms. The study suggests how the combination of autonomous technologies can accelerate genetic modification. By performing the first experiments with a programmed robotic system that can process liquids with remote control, transgenic bacterial or human cells that can express green fluorescent protein were created by performing genetic transfer in bacteria or human cells in an automated manner.

With the CRISPR.BOT project, for the first time, genetic modifications were carried out in cells with an autonomous approach, without human intervention, and subcloning processes were achieved by recombinant viruses encoding CRISPR gene engineering. This smart and versatile robotic system suggests a cost-effective approach for molecular biology laboratories and also contributes to laboratory functions with a wide range of applications.

## Materials & Methods

For molecular biology experiments with the CRISPR.BOT, we first prepared plasmid DNA, and GFP or CRISPR-Cas9/gRNA encoding lentivirus with the following methods.

### Transformation and Isolation

Competent *E. coli* DH5α strain (NEB® 5-alpha Competent *E. coli* (High Efficiency) NEB C2987H) were cultured in a liquid LB medium at 37°C. The bacterial strain was stored at -80°C as stock in LB medium with added glycerol. Bacteria were transformed with the plasmid DNAs to be used in the CRISPR.BOT studies using the heat shock method according to the manufacturer’s instructions. Bacteria were inoculated on LB agar plates with ampicillin and incubated at 37°C overnight. Plasmid isolation was performed from selected bacterial colonies with the CompactPrep Plasmid Maxi Kit (#12863, QIAGEN). DNA concentration was measured in ng/µl with Microplate ELISA Reader (FLUOstar Omega) and it was evaluated whether the purity value was between 1.8<A260/A280<2.0.

### Lentivirus Production

The previously designed 3 gRNA sequences were synthesized with the pU6 promoter by GenScript and cloned into the lentiviral pHIV-EGFP (Addgene, #21373) plasmid (Odabas et al., 2022). For lentivirus production, CRISPR-Prime Editing system encoding lentiviral pLenti-PE2-BSD plasmid DNAs (Addgene, # 161514) encoding CRISPR-PE containing the blasticidin resistance gene, control pHIV-EGFP non-coding gRNA1/2/3, and coding GFP and pHIV EGFP coding pU6-gRNA1/2/3 and GFP were transfected with pSPAX2 and pVSV-G envelope and packaging plasmid DNAs. After all plasmids (2:1 pSPAX2: 1 pVSV-G) were treated with P-PEI (Polyethylenimine) transfection reagent (Merten et al., 2016), lentivirus production was performed using the host cells, HEK293T (Odabas et al., 2022). Packed recombinant lentiviruses were obtained by collecting HEK293T cell supernatants 72 hours after transfection. Produced lentiviruses were concentrated 100x with Lenti-X Concentrator (#631232, Takara) to increase virus concentration. For the titration of lentiviruses, titration steps using Jurkat (RPMI medium containing 10% fetal bovine serum, 200 U/ml penicillin/streptomycin antibiotic, 2 mM L-glutamine, 1X MEM vitamin solution, and 1X NEAA) cells in a previously published study were performed (Taştan et al., 2020). After the cells were transferred to 96-well plates, Jurkat cells containing pLenti-PE2-BSD (Addgene, #161514) were treated with 10 µg/ml blasticidin for selection of blasticidin and incubated for 72 hours. Afterwards, cell viability was tested by staining with a 7-AAD antibody in flow cytometry. Finally, cells encoding CRISPR-PE with >99% blasticidin resistance were selected and cultured. Titration of gRNA-GFP and control GFP lentiviruses was calculated by measuring the GFP expression level in flow cytometry. Subsequently, using CRISPR.BOT, 3 gRNA-GFP or control GFP lentivirus were individually transduced into CRISPR-PE (+) Jurkat cells.

### Cell Survival Analysis

Survival analysis was performed to control the cell viability of Jurkat cells. Trypan blue (Biological Industries, #03-102-1B) was applied to identify and count surviving cells. Cell counting and viability analysis were performed with the BIO-RAD TC20 Automated Cell Counter.

### Statistical Approach

Data recorded by flow cytometry were analyzed using CytExpert (Beckman Coulter). Statistical analyses were performed using GraphPad Prism 8.0.1 software (GraphPad Software, Inc., San Diego, CA, USA) with a two-tailed t-test using independent mean values. Error bars represent Standard Error of Means (SEM). For all experiments, significance was defined as *p<0.05 and *NS*= Non-Significant.

## Results

### Construction of CRISPR.BOT autonomous and remotely controllable robotic system

The LEGO Digital Designer application was used to construct the CRISPR.BOT v1 robotic system with LEGO Mindstorms Education EV3 and the first entire installation benefitted from the study published by Gerber LC et al. in 2017. However, CRISPR.BOT v2 installation stages were completed from the beginning using the LEGO Digital Designer program as reported in the literature (Gerber LC et al., 2017). To watch the construction of CRISPR.BOT version 1 (V1): https://drive.google.com/file/d/1dAMVvnkI0DES3bo71ki12z7p-iDurkRc/view?usp=sharing

The rail has been enlarged to place the plates, petri dishes, and cuvettes that were used in the experiments planned to be carried out in a more robust and wider area (**Figure 2A**). A wide area was created by increasing the rail system from one row to three rows (**Figure 2B**). There is one color sensor in the CRISPR.BOT v1 robotic system. This color sensor in the LEGO Mindstorms EV3 set; can detect 8 different colors and measure the values of ambient light or reflected light. The primary purpose of the experiments with the color sensor was to ensure that the color sensor stops at the desired colors. By providing the color sensor integration with the syringe, the colored papers on the cuvettes stopped with the detection of the color sensor, and the syringe system provided the desired movement. Colored papers were stuck in the middle of the tubs for the color sensor to recognize (**Figure 2B**).

**Figure 1.**
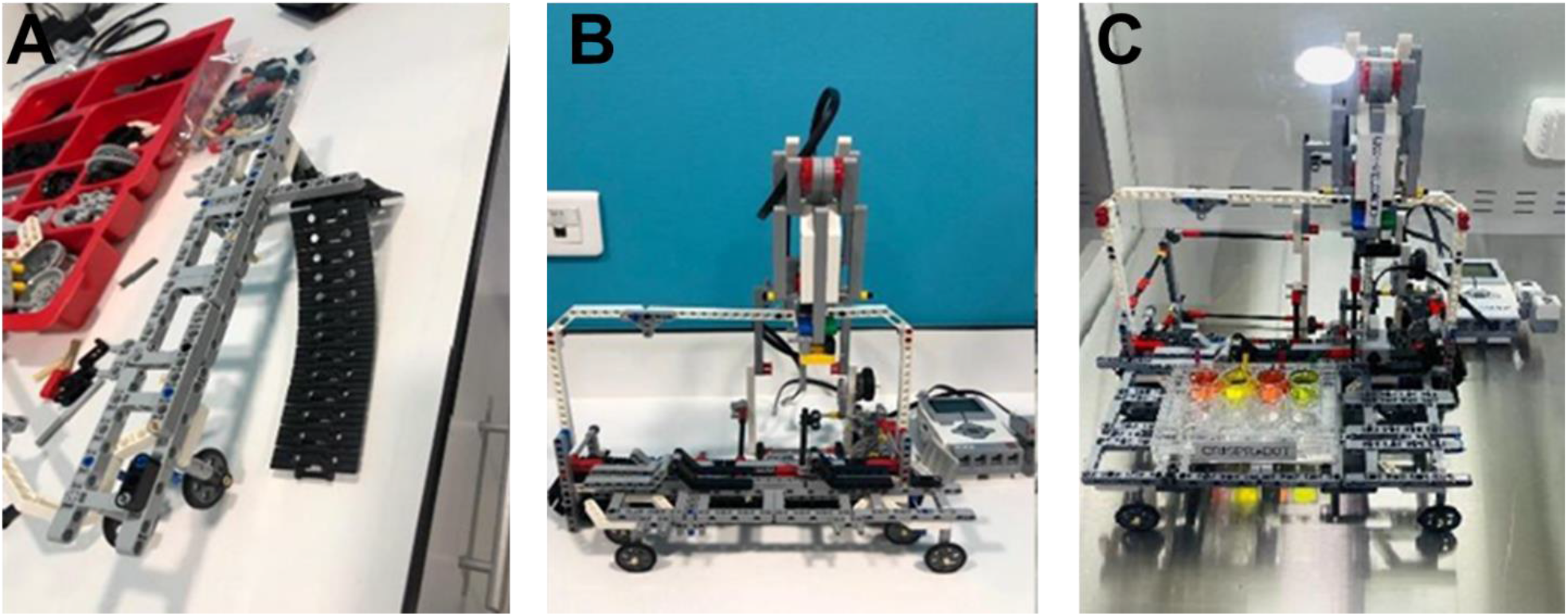
CRISPR.BOT V1 setup steps. **A**. Installation of the rail system for CRISPR.BOT V1. **B**. The assembled version of the V1 robotic system using the LEGO Digital Designer program and the spare parts in the LEGO Mindstorms set. **C**. By expanding the rail system, the plates to be used in the experiments planned to be carried out more comfortably.

**Figure 2.**
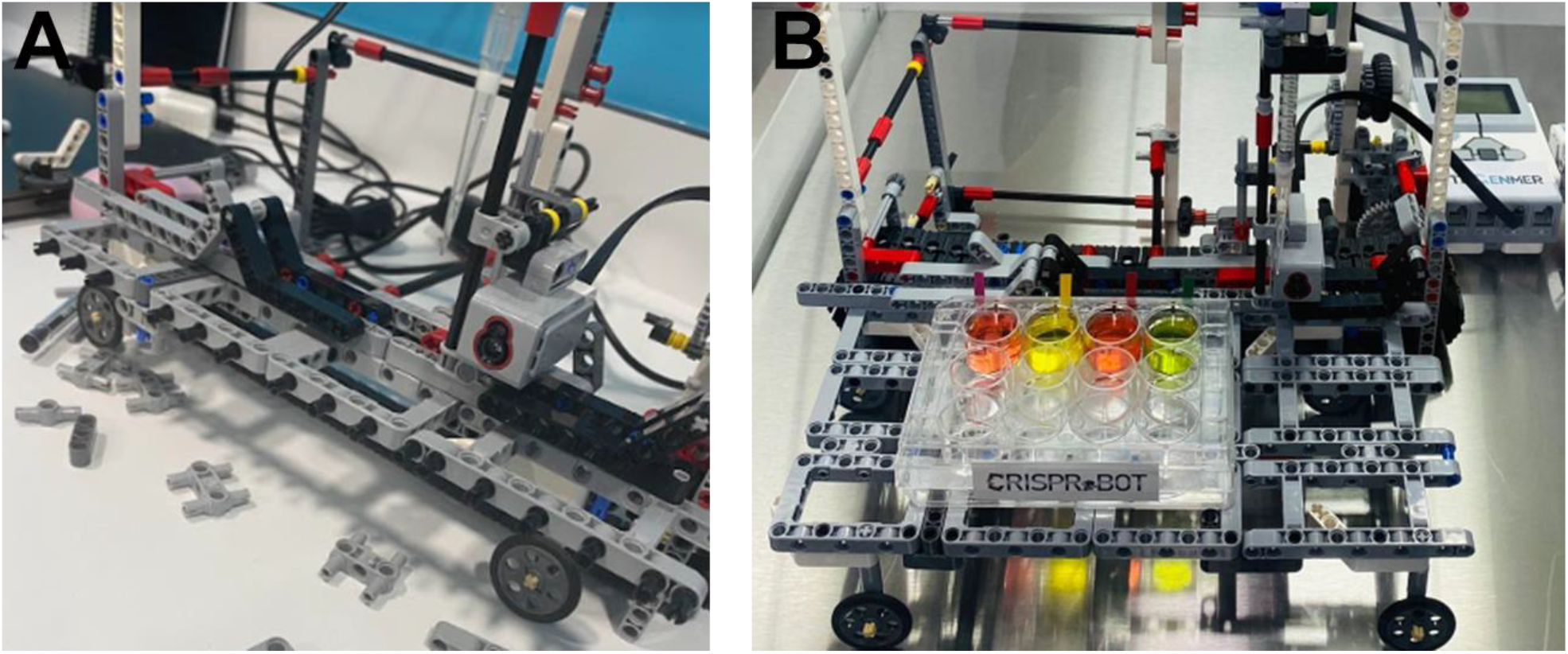
Rail system. **A**. Expansion of the rail system. **B**. Colored papers integrated into the plate for the plate carrier rail system and color sensor.

For the V1 robotic system, the syringe plunger was cut by the system in the robot and fixed to the LEGO piece with the help of glue and connected to the syringe (**Figure 3**).

**Figure 3.**
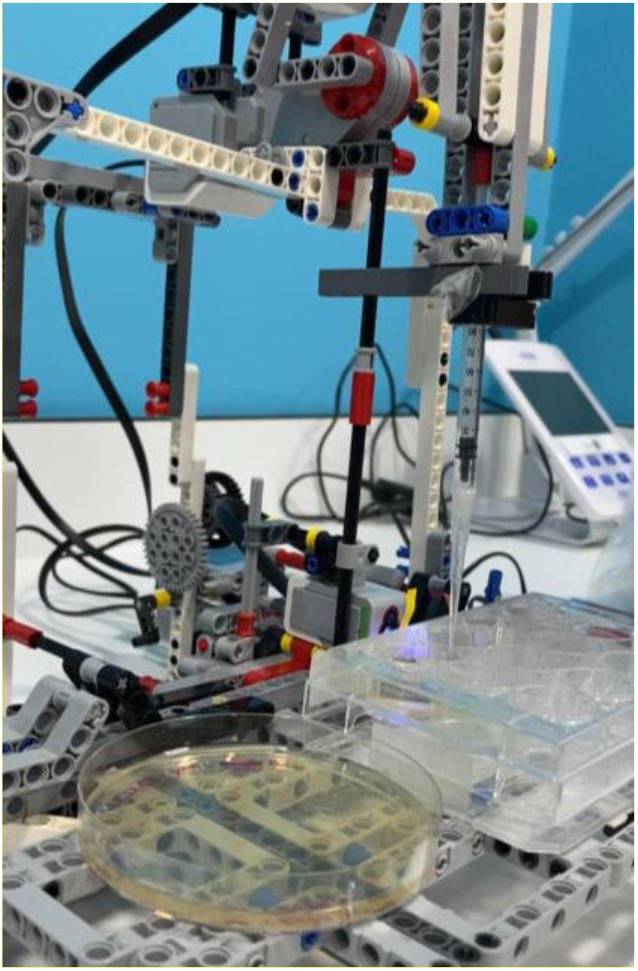
The position of the syringe on the cuvette.

Certain blocks have been prepared for the following experiments by downloading the program. The purpose of these blocks is to perform the commands in the blocks, respectively, when each block represents a movement and is added below the command that initiates the movement (**Figure 4**).

**Figure 4.**
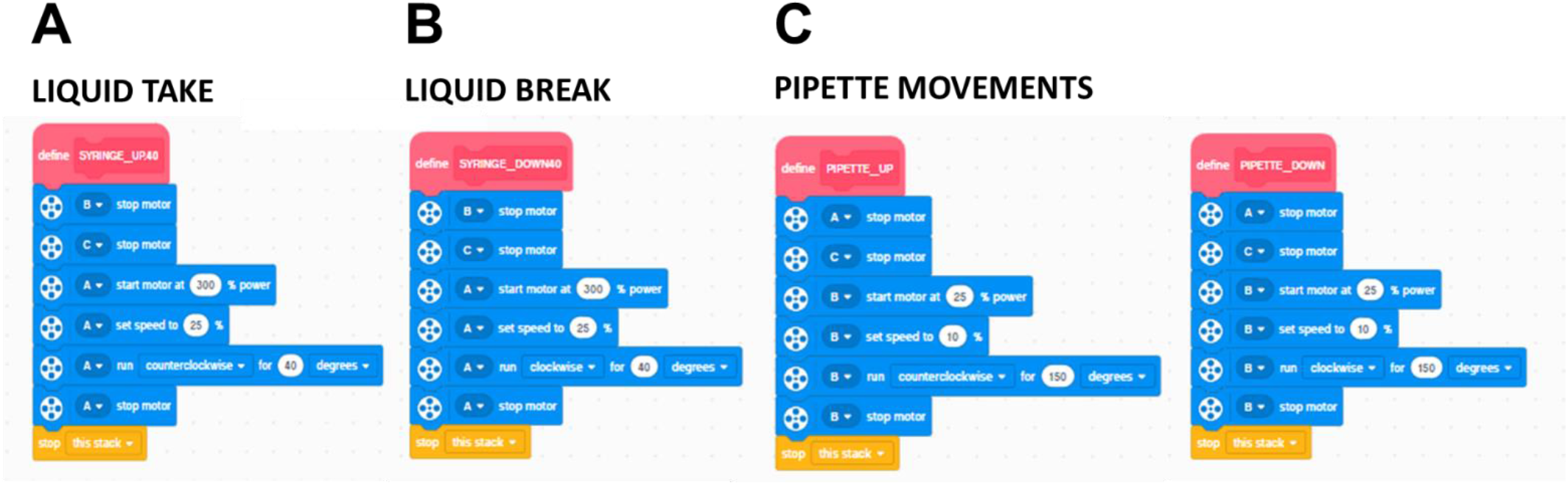
Blocks used in experiment programming. **A**. Liquid Take: It fulfills the liquid intake task of the pipette system. With the 40° movement of the large servo motor that provides the movement of the pipette system, the piston connected to the syringe goes up and takes liquid. **B**. Liquid Break: It performs the liquid delivery task of the pipette system. It provides the opposite action of the fluid intake process and leaves the fluid taken back. **C**. Pipette Movements: They are the blocks that perform the up and down movements of the pipette system. The power and speed are kept low so that the liquid enters slowly and precisely.

Every movement of the robot is coded to process liquids in microliter volume with velocity and angular measurements (**Figures 5 and 6**). With the degree reset block, it was aimed to reset the calibration of the robot before each experiment and to perform its movements more accurately. To ensure that the syringe draws the same amount each time in the experiments to be carried out, different experiments were made on the programming and the liquids were measured with the help of a pipette after each liquid draw. As can be seen from **figure 5**, because of the velocity-microliter measurements, it was observed that the velocity change did not change the measured microliter amount.

**Figure 5.**
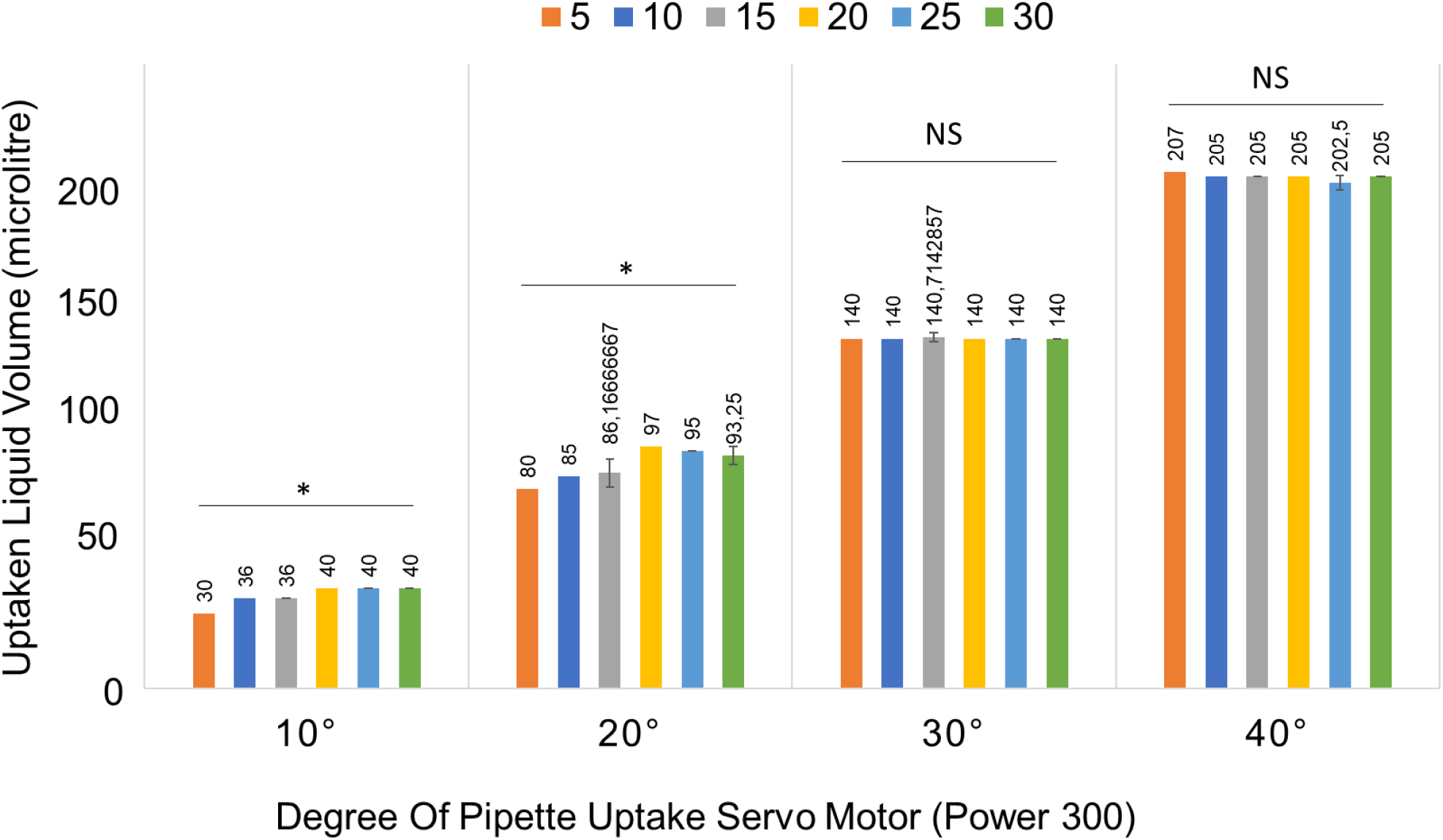
Power constant (=300), speed-microliter table. At a constant power of 300, microliter measurements were quantified at different speeds of Liquid Take, Liquid Brake, and Pipette Movements for 10°, 20°, 30°, and 40°, and it was seen that there was not much difference in the withdrawn microliter even though the speed was increased.

**Figure 6.**
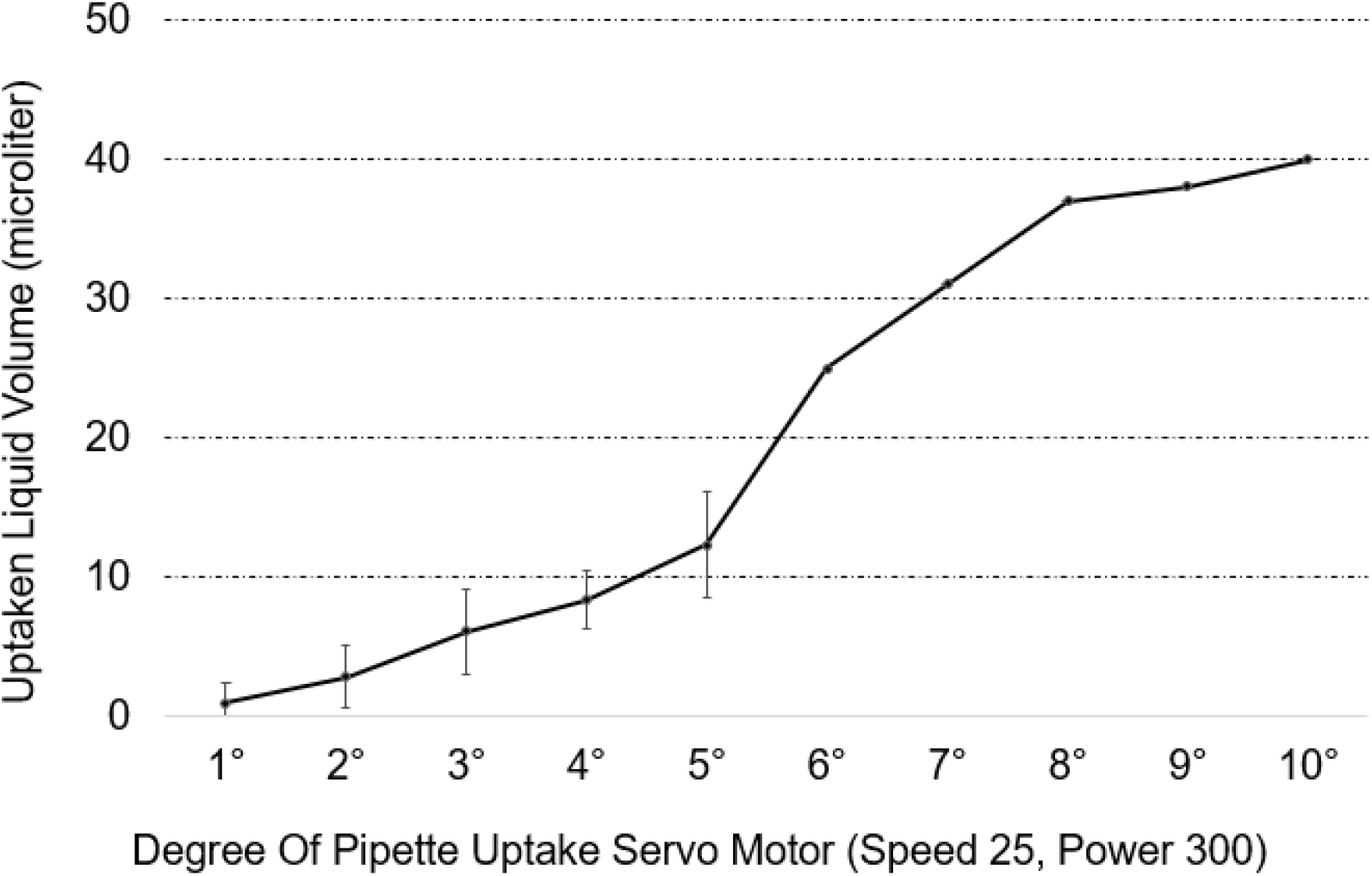
A graph of utilized microliters with different Servo Motor uptake angles including 1°-10° (Constant Pippette clock speed (=25) and power (=300)).

As a result of the measurements, it is seen that the CRISPR.BOT robotic system can draw low microliter liquids between 1°-10° and 0-40, but there are deviations in the liquid withdrawals between 1°-5° **(Figure 6)**.

After many trials and finding the right programming, experiments were carried out at different speeds and at different angles to understand the connection between velocity and measurement. Also, a chart was drawn up for each and it was determined that the different velocities do not affect the volume to any constant degree (**Figures 5 and 6**). By determining which angle to use in experimental programming from the graphics, its programming was arranged, and the first bacterial transformation experiment was carried out with the robot.

For the new version installed, the system mechanism has been completely changed and new pipette movements have been added to the robotic system. While the pipette mechanism can only move up and down in CRISPR.BOT V1, right and left movements can be provided in the installed CRISPR.BOT V2. With a rail system under the CRISPR.BOT V1 pipette, movement is provided and the well on the well plate to be processed is brought to the level of the pipette, while with CRISPR.BOT V2, the pipette can be taken to the desired location with its rail system. The kits in the LEGO Mindstorms series have the software and necessary hardware to create a programmable customizable robotic system (Montes, 2021). The LEGO Mindstorms EV3 Home Edition software was downloaded from the LEGO website and certain functions were added to the robots (**Figure 4**). This study aimed to test the experimental approaches by providing the robot with the ability to process liquids with the help of LEGO Digital Designer and LEGO software. (Gerber, 2017; Scaradozzi,2020). The wheel at the top left in the image is connected to a servo motor and alone performs the task of pulling and releasing liquid, which we call the syringe system and the mechanism linked to **Fig 7A**. While **Fig7A** can only pull and release fluid by fixing the syringe plunger, when placed inside **Fig 7B** and connections are made, it has two more motion systems. The first of these provides up and down movements of the pipette tip so that it can enter the liquid. On the other hand, the wheels at the four corners of the mechanism act as a wheel and move on the gears and can move on the robotic system by being placed on the system on the frame.

**Figure 7:**
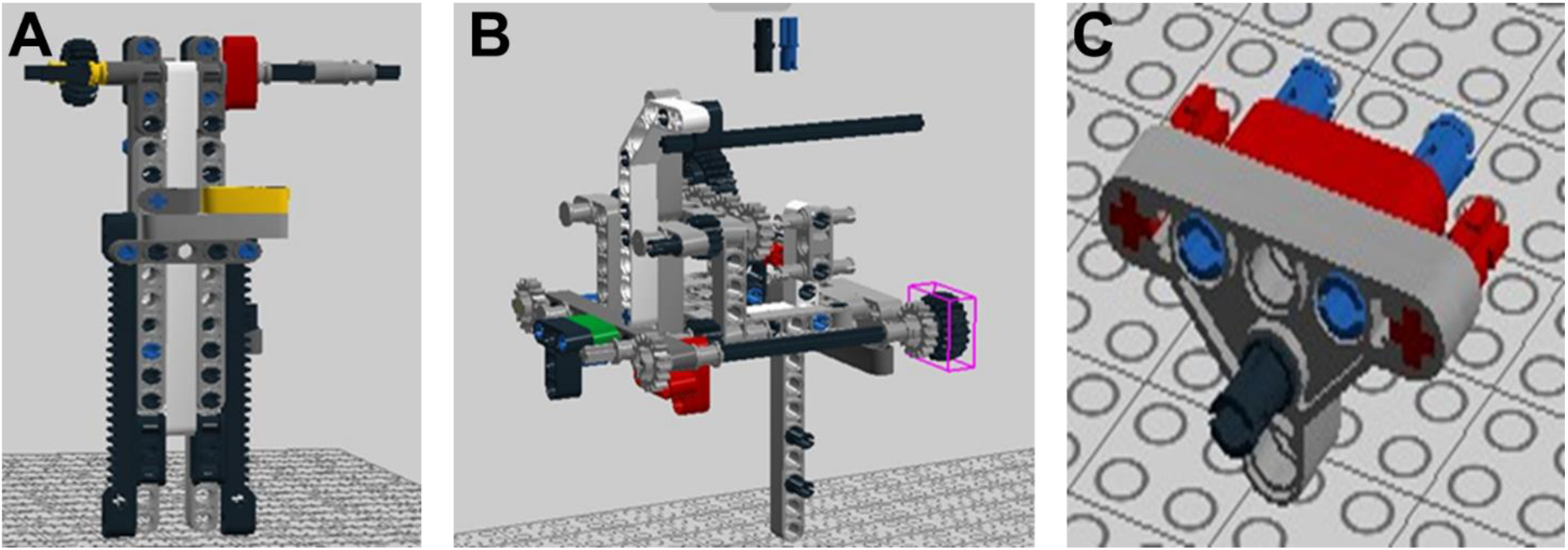
Construction of the CRISPR.BOT V2 pipetting system. **A**. It is the mechanism that provides the movement of the syringe and is placed in the middle part of **B**. The movement of the wheels is provided by toothed rail plates and the up and down movement of the straw is provided. **C**. It is a part that keeps the syringe connected to the system.

To provide the piston movement connected to the syringe, the piston must be connected to a LEGO piece and combined with the wheels and provide up and down movement (**Figure 8**).

**Figure 8.**
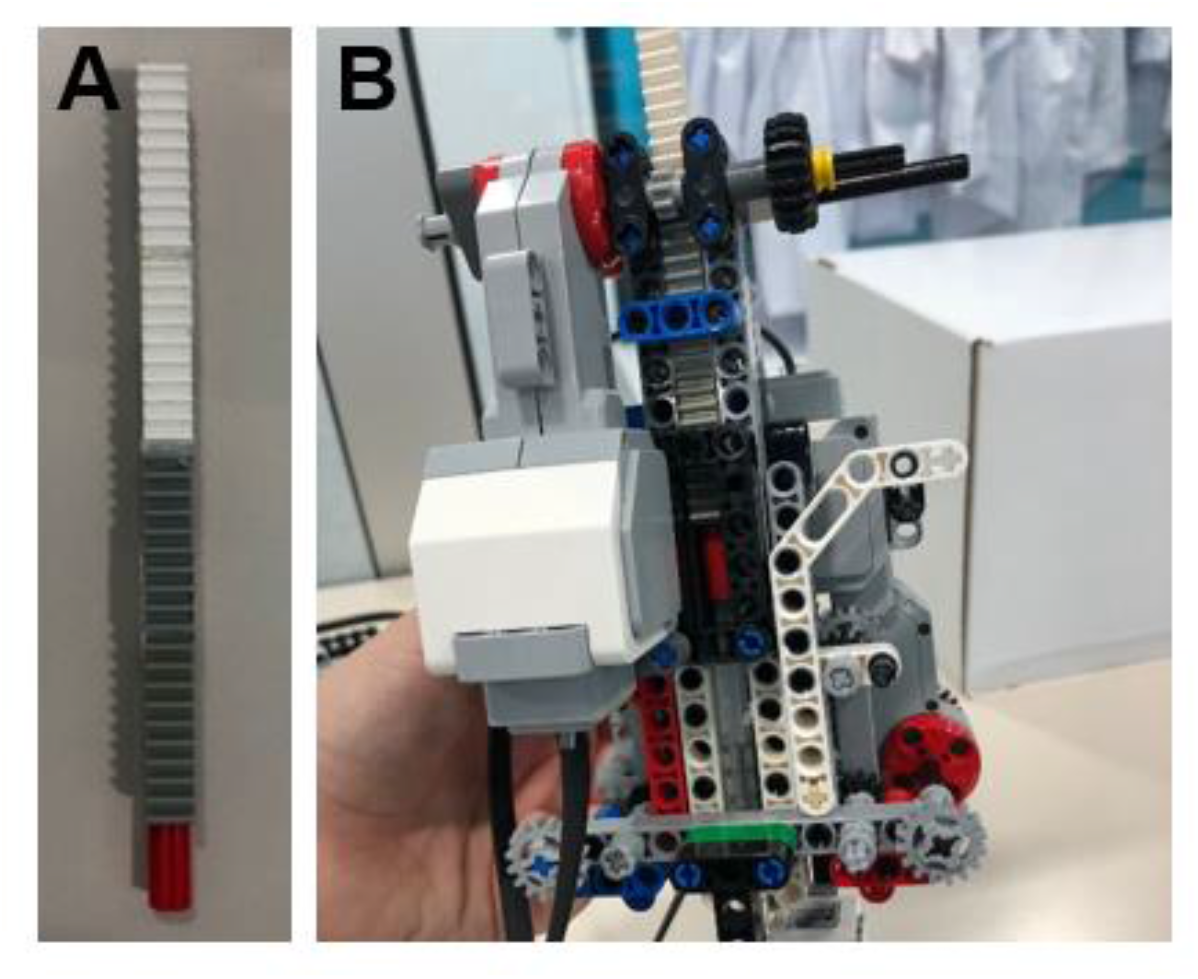
Alternative parts are used to provide the syringe movement connected to the plunger. **A**. By placing the syringe plunger inside the red part, the movement of the part connected to the piston is provided by the movement of the impellers connected to the servo motor. With this system, the movement of the piston up and down is provided, and the fluid exchange process is carried out. **B**. The prepared mechanism is placed in the pipette system.

In the CRISPR.BOT V2 robotic system, there are 3 different motors integrated into the pipette module. Two of them are large servo motors and one is a medium servo motor. One of the large servo motors provides the movement of the robot on the frame. Another large servo motor provides the movement of the pipette system, which provides the liquid exchange process (**Figure 9A**). With the operation of the medium servo motor, the movement of the impellers allows the pipette, which is integrated into a rail mechanism, to enter the liquid, providing up and down movements (**Figure 9B**).

**Figure 9.**
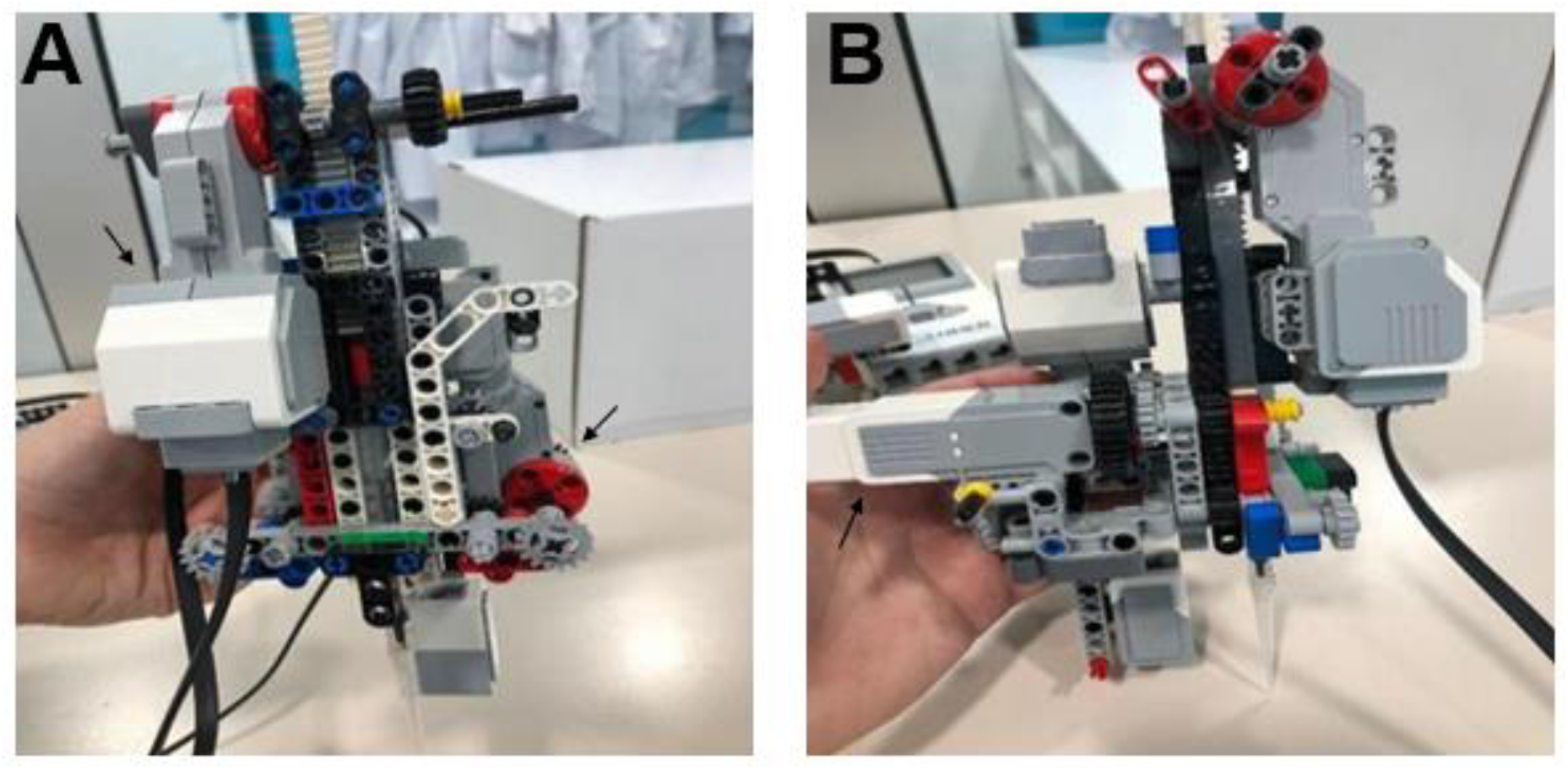
Integration of the CRISPR.BOT pipetting system into the servo motor. **A**. The movement of the impeller connected to the large servo motor on the rail plate provides the movement of the pipette system, which performs the liquid exchange process. **B**. Side view of the pipette system. The up-down movements of the pipette system are provided by the medium servo motor moving the wheels.

The movement of the wheels is provided by the toothed rail plates and the upward and downward movement of the straw is provided (**Figure 10**).

**Figure 10.**
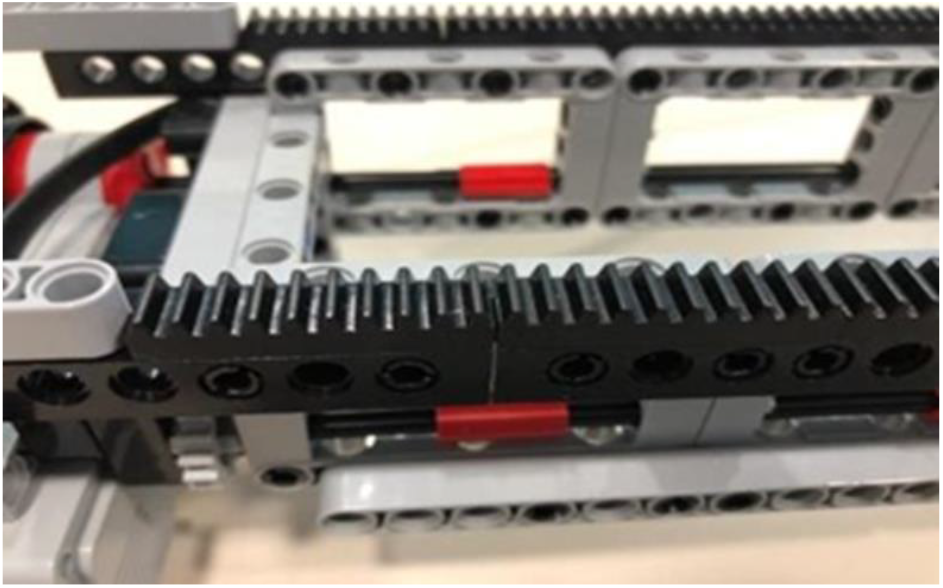
The gear piece enables the wheels to move on it.

The pipette system, which performs pipette movements and liquid exchange operations, is placed on the frame. The movements are carried out by the gear parts on the frame and the wheels at the bottom of the pipette system. The frame is fixed and does not have any mobility (**Figure 11**).

**Figure 11.**
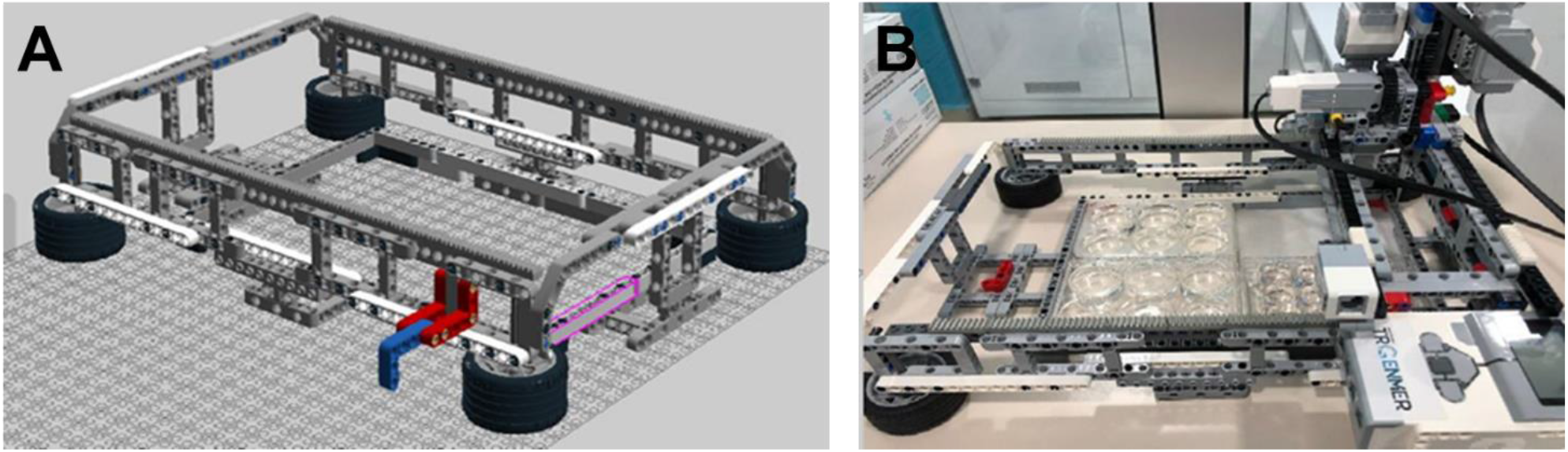
CRISPR.BOT is the bottom part of the robotic system, called the frame. **A**. Image of the frame setup made from the LEGO Digital Designer program. **B**. It is the bottom part of the robotic system, called the frame. The pipette system, which performs pipette movements and liquid exchange operations, is placed on the frame. Movements take place thanks to the toothed parts on the frame and the wheels at the bottom of the pipette system. The frame is fixed and does not have any mobility.

It is the mechanism that provides the movement of the pipette system to the right and left and back and forth movements on the frame (**Figure 12**).

**Figure 12.**
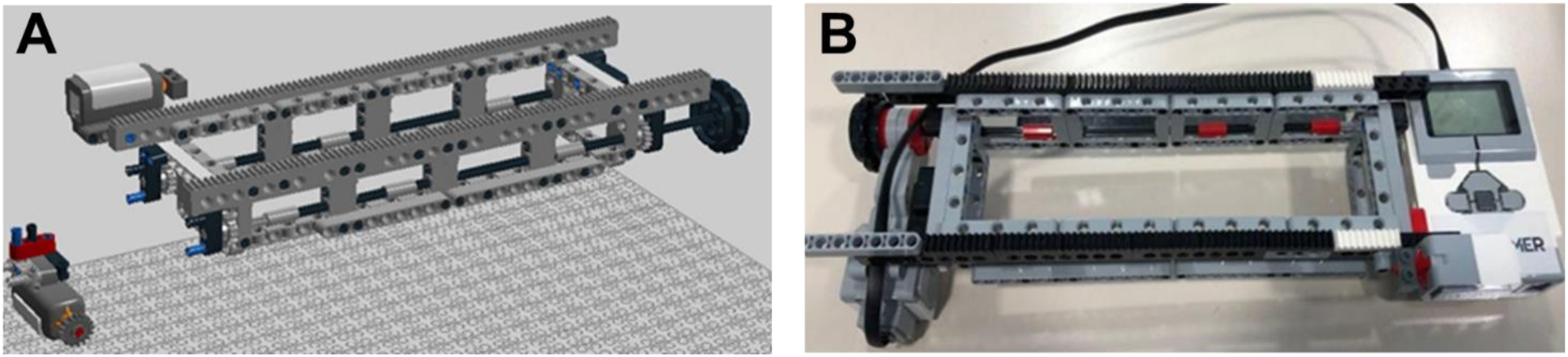
The system provides the liquid transition movements of the pipette between the other wells. **A**. The mechanism that provides the movement of the pipette system on the frame, was designed with the LEGO Digital Designer program. **B**. It is the mechanism that enables the pipette system to move right, left, forward, and backward on the frame. The pipette provides fluid passage movements between other wells.

For the new version that was installed, the system mechanism was completely changed, and new pipette movements were added to the robotic system. While the pipette mechanism provided only up and down movements in the previous version, right and left movements can be provided in the new version that has been installed. In the previous version, while the well on the well plate to be processed was brought to the level of the pipette by a rail system under the pipette, with this version, the pipette can be taken to the desired location with its rail system (**Figure 13**).

**Figure 13.**
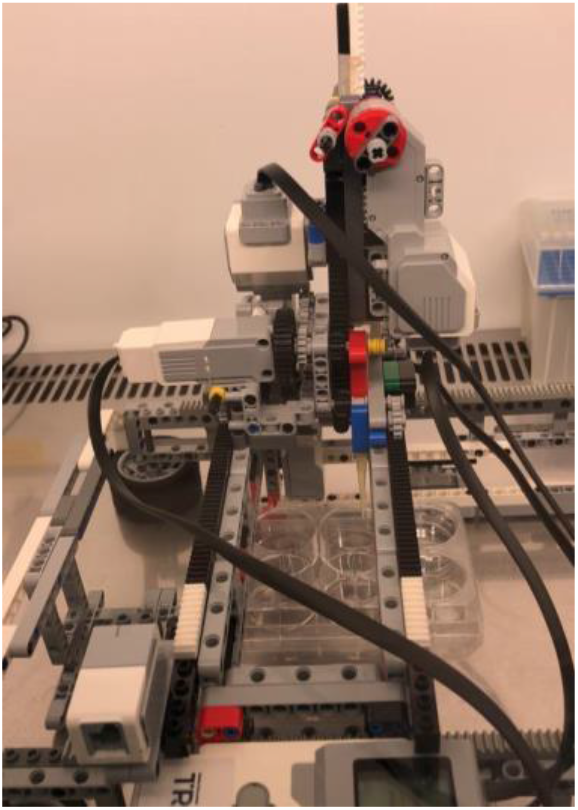
Pipette system that provides the right and left movements of the pipette.

In the speed measurement optimization phase of the CRISPR.BOT V1 version, many trials were made on power and speed in its programming, and since it was seen that the use of high power was more effective, the optimization studies of CRISPR.BOT V2 were continued over this and the power and speed were kept constant at high degrees. For the 6 well plate, 12 well plate and 96 well plate planned to be used in the experiments, separate plans were made, and their programming was prepared. First, planning was made by dividing the rail movement into two. The first stage is the rail movement of the mechanism that takes the straw on the frame to the desired position, that is, the back-and-forth movements. In this, separate calculations were performed for each well plate.

### Genetic Transformation in Bacteria

Based on the velocity-microliter measurement charts prepared for the targeted bacterial transformation experiment, the degrees required to carry out the experiment without error by drawing the required amount of liquid were determined. To prepare for the experimental programming, the bacterial transformation test stages to be carried out were schematized and planned (**Figure 14**). To watch the transformation experiment: https://drive.google.com/file/d/1UzVVautbBzsdWbKYAVo2E3aOgvmZmMPE/view?usp=sharing

**Figure 14.**
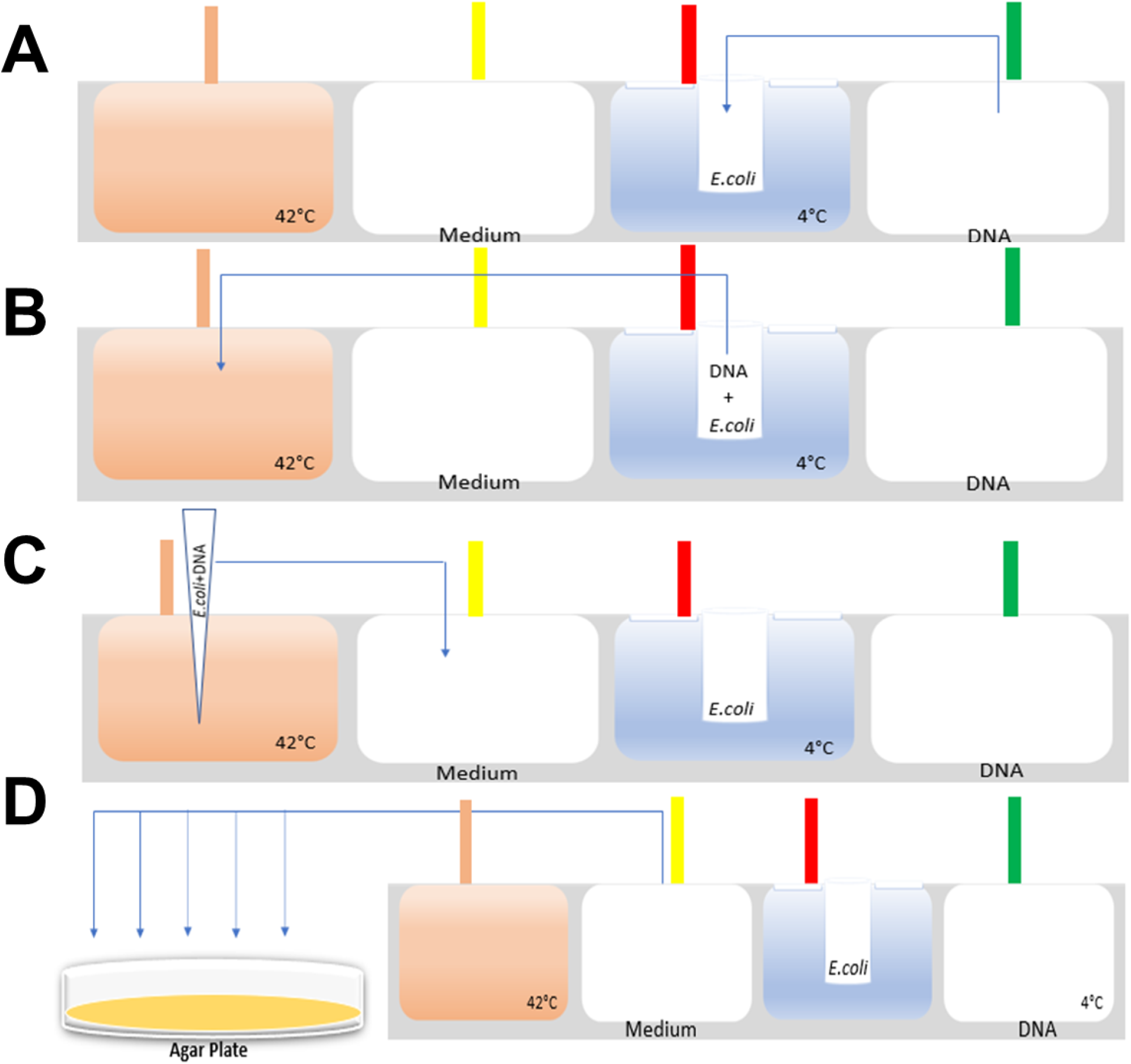
The schematization of bacterial transformation experiment steps. **A**. Take 10-20 µl of DNA at an angle of 5° and add 50 µl of *E*.*coli* in the transwell at 4°C. Then wait 30 minutes. **B**. Take 60 µl of DNA + *E*.*coli* at an angle of 15° and wait for 20-30 seconds by immersing the pipette in 42°C hot water. **C**. Remove the pipette, which is immersed in 42°C hot water, and leave *E*.*coli*+ DNA in an 800 µl medium. Then shaking is done 3 times at intervals of 5 minutes for 15 minutes. **D**. Take 205 µl 3 times at an angle of 40°, leave it in different parts of the petri dish and allow it to spread by shaking for 10 seconds.

White colonies formed as a result of DNA transformation with an autonomous remotely controlled robotic system were determined in red and blue circles (**Figure 15**).

**Figure 15.**
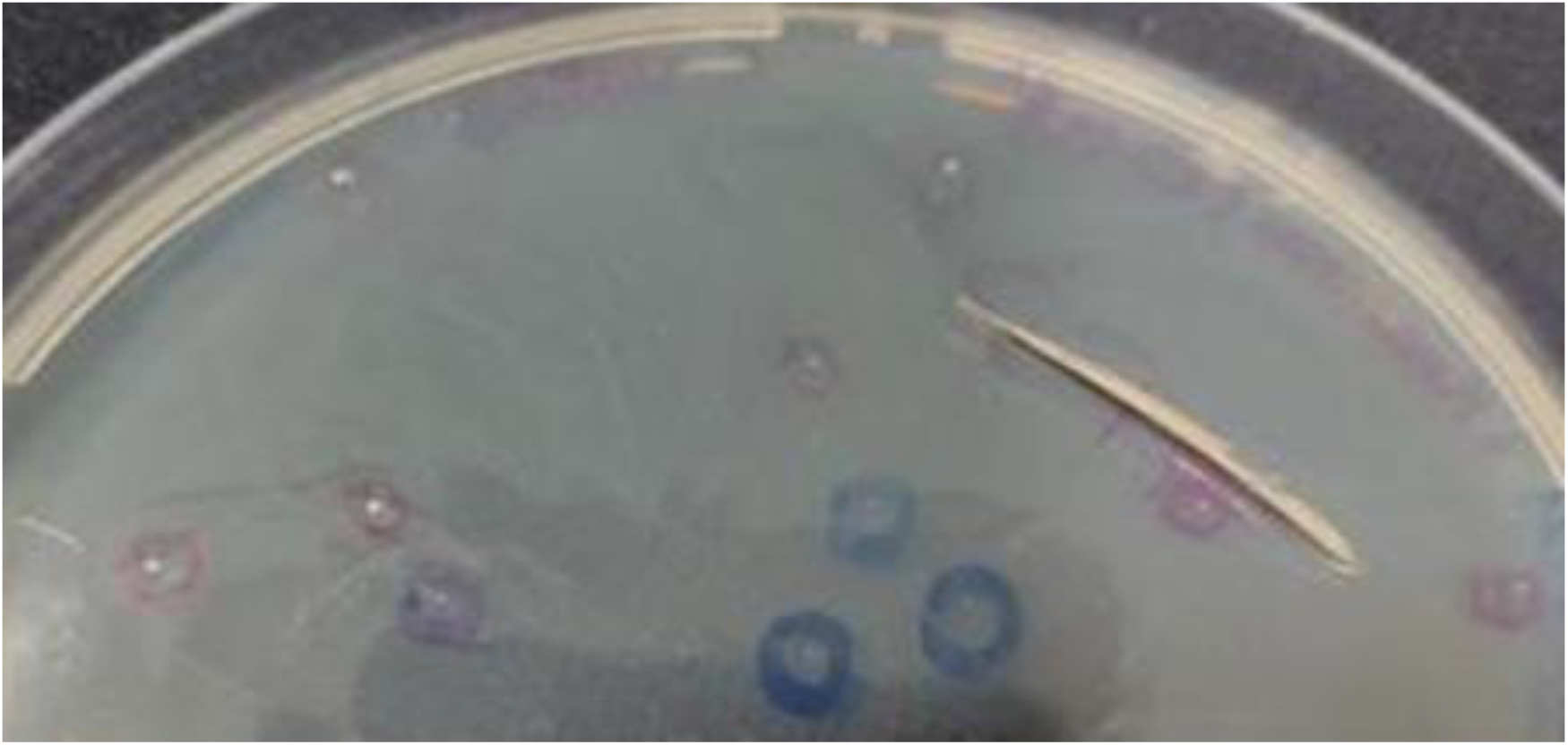
Bacterial Colonies After Transformation using CRISPR.BOT. Transgenic white bacterial colonies formed as a result of the transformation of plasmid DNA encoding GFP and Ampicillin (Amp) resistance gene into *E*.*coli* bacteria.

#### Genetic Transfer in Human Cells

With the 2nd version of the robotic system, the first experiments were carried out to prove that human cells can be genetically modified with viruses. The experiment was carried out; It was aimed to prove that GFP lentivirus can genetically modify human cells with an automated system (**Figure 16**). In the experiment planning, appropriate coding was performed for the 4 wells of the 12-well plate, side by side, with the first well as a control and increasing amounts of GFP lentivirus transfer to the other three wells (**Figure 17**). Cells were seeded in 4 wells of the 12 well plate to be used first, with 2,5×10^4^ Human Jurkat cell lines in each well, and each well was completed with RPMI medium with a total volume of 500 µl. Plates were placed in the frame of the robotic system by adding 1 ml of GFP encoding recombinant lentivirus to one well of the other 12 well plate. For watching: https://drive.google.com/file/d/1L4GX1Z_ssDuZoal-4KZIC-ly8b6oh63S/view?usp=sharing

**Figure 16.**
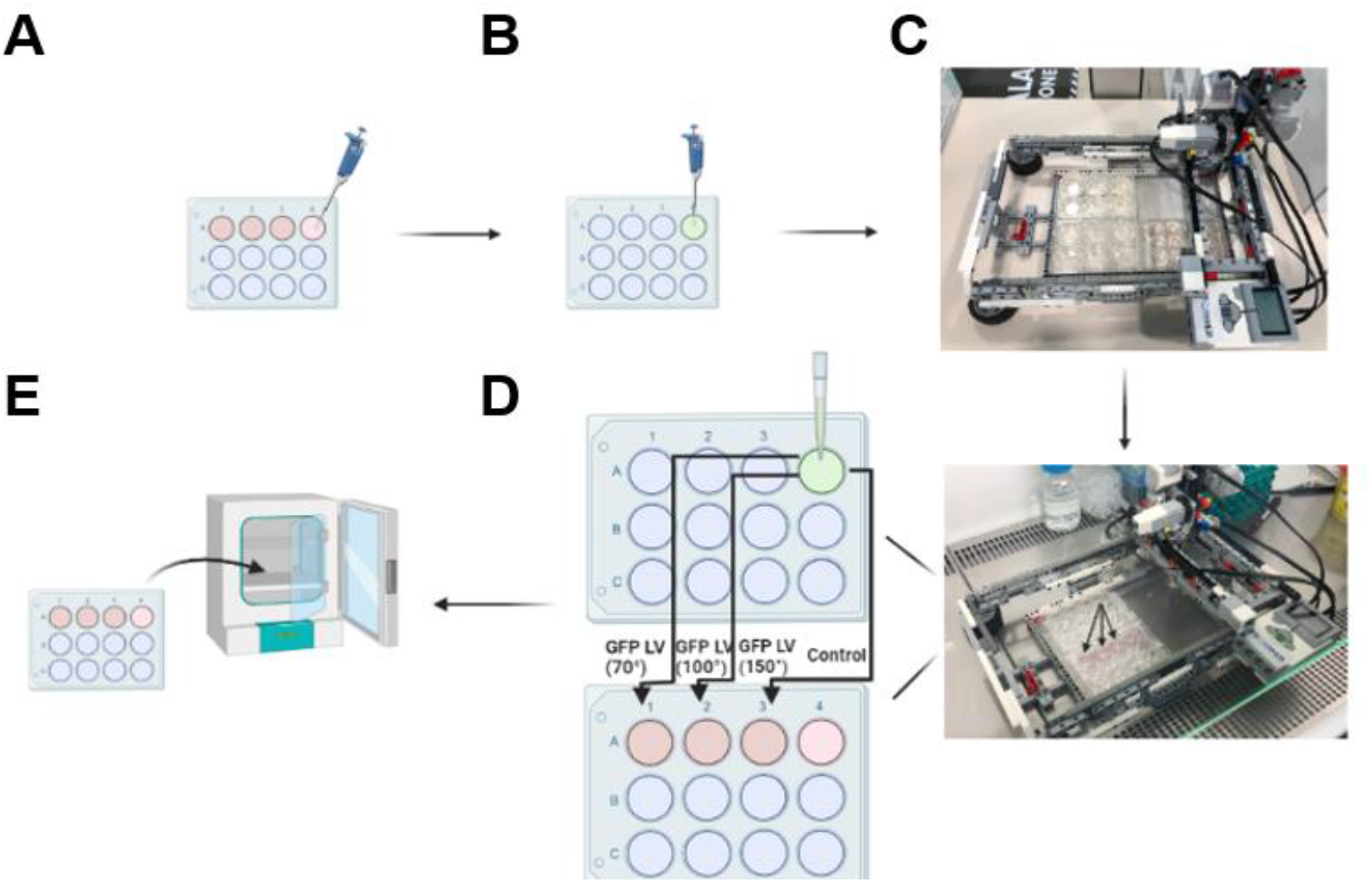
Genetic Transfer in Human Cells. **A**. Performing Cell Sowing, Jurkat Cells were cultivated in 4 wells at 2.5×10^4^ cells/well. It was cultured in RPMI medium with a total volume of 500μl. **B**. Virus Sample Preparation, 1 ml of GFP (Green Fluorescent Protein) Lentivirus was placed in one well of the 12 well plate. **C**. Integration of Plates into the Autonomous Robot System, 12-well plates were placed in the frame of the Robot. **D**. Addition of GFP Lentivirus to Cells, 70°, 100°, and 150° extractions were made from the well containing GFP Lentivirus (GFP LV) in a 12-well plate and added into wells 1, 2, and 3. **E**. Incubation, after the robot performed the experiment, the 12-Well plate was placed in the incubator.

**Figure 17.**
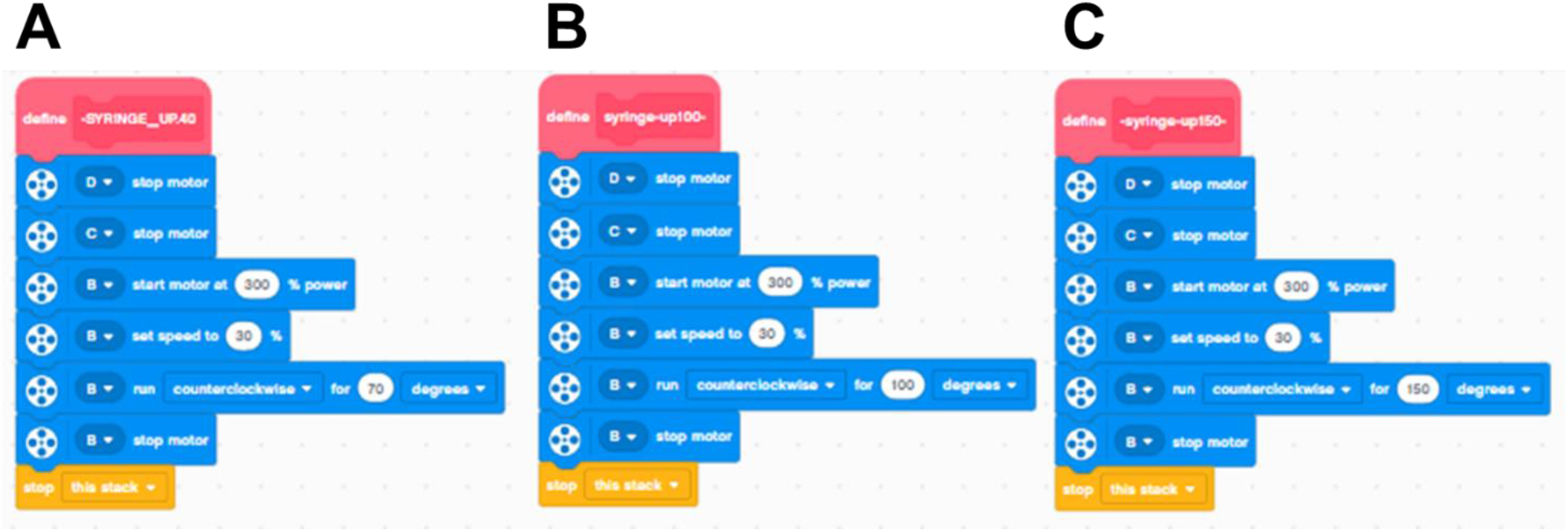
70°, 100°, and 150° blocks were used for Genetic Transfer in Human Cells experiment programming. **A**. 70°, **B**.100°, and **C**. 150° blocks that activate the pipette system are prepared in the experiment programming and perform the liquid withdrawal process.

The robot performed GFP lentivirus extraction at a 70 ° angle (60µl-80µl) and added the virus to the 1st well in which Jurkat cells were seeded in the first 12-well plate (**Figure 18**). Then, the CRISPR.BOT returned and added GFP lentivirus with a 100 ° angle (90µl-120µl) and added the virus to the second well in which Jurkat cell was cultivated. Finally, the robot added the virus to the 3rd well by shooting with a 150° angle (160µl) from the same virus. As a control, RPMI medium was added to the 4th well at an angle of 100 degrees. The plate was placed in a cell culture incubator at 37 C containing 5% CO2. For the control of cell number and viability, cells were analyzed with the trypan blue and a Cell Counting device (Biorad TC20) (**Table-1**). This shows that CRISPR.BOT does not have cytotoxicity on the cells.

**Figure 18.**
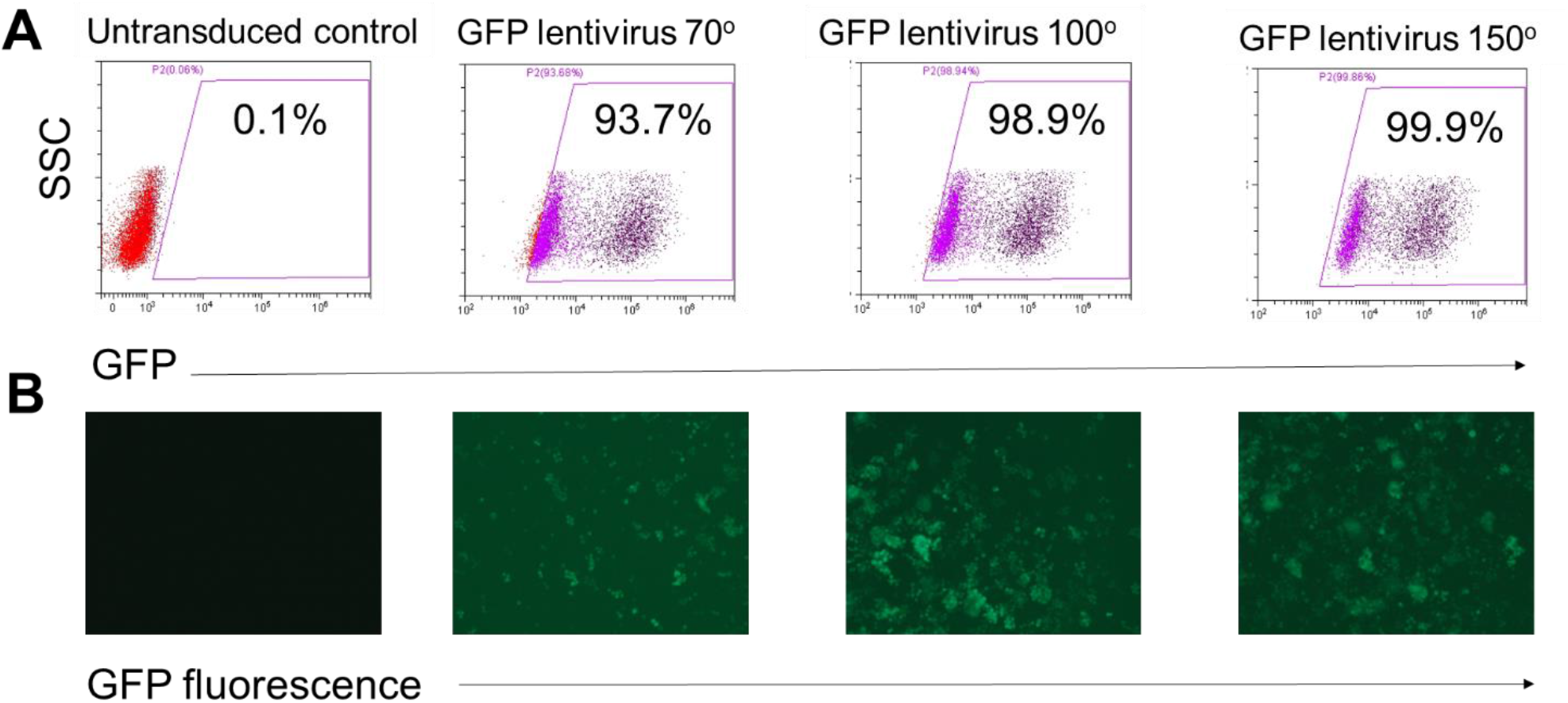
Analysis of GFP protein expression in CRISPR.BOT modified transgenic Jurkat cell line using **A**. flow cytometry and **B**. fluorescence microscopy at 4X.

**Table 1:** Cell Count and Viability Rates of Jurkat cells After 72 Hours of Transduction

| Conditions | Viable cells | Viability |
| --- | --- | --- |
| Control untransduced cell | 2,04x10<sup>5</sup> | 97% |
| 70° GFP Lentivirus | 2,53 x10<sup>5</sup> | 84% |
| 100° GFP Lentivirus | 3,19x10<sup>5</sup> | 94% |
| 150° GFP Lentivirus | 2,15x10<sup>5</sup> | 78% |

With this experiment, it was found that there was no GFP expression in the control group, but an increased GFP expression at 70-100-150-degree angles, respectively.

GFP expression levels were determined 72 hours after experimenting by Flow Cytometry analysis and fluorescent microscope imaging (**Figure 18**).

As a result of the experiment, it was observed that there was no virus expression in the control group, and an increased virus expression was observed in the injection process made by the robot with the pipette system at 70°-100°-150° angles (**Figure 18**). Thus, it has been proven that the genetics of cells can be changed with a cost-effective autonomous system with viruses transferred in different amounts. The experiment aimed to automatically produce transgenic cells and work remotely with autonomous systems without touching the viruses, which are dangerous to study. The results obtained from the experiments carried out, it is the prototype of a system in which viruses can be produced and used without human touch, and vaccine and drug studies can be carried out in studies to be carried out with viruses such as SARS-CoV2 which is infectious.

### CRISPR Gene Modification

With this experiment, it was aimed to transfer three different CRISPR guide RNAs of the robot to the cells. The main purpose of the evidential experiment for the future is to carry out the experiment by understanding which guide RNA should be added to which cell with the CRISPR system to perform the required genetic modification in the cell. Cells were seeded in 4 wells of the first 12 well plate with 5×10^4^ Human Jurkat cells in each well, and the total volume was completed with RPMI medium to be 500 µl in each well. Lentiviruses encoding guide RNA 1 (gRNA1), guide RNA 2 (gRNA2), and guide RNA 3 (gRNA3) were added to the wells of the other 12 well plates, respectively, by CRISPR system, diluted 1:10 in RPMI medium. The plate was lifted into a cell culture incubator containing 37^0^C 5% CO2 (**Figure 19**). For watching: https://drive.google.com/file/d/16vGq9T99VS1300GIFbGGO1EZfKEtbSsy/view?usp=sharing

**Figure 19.**
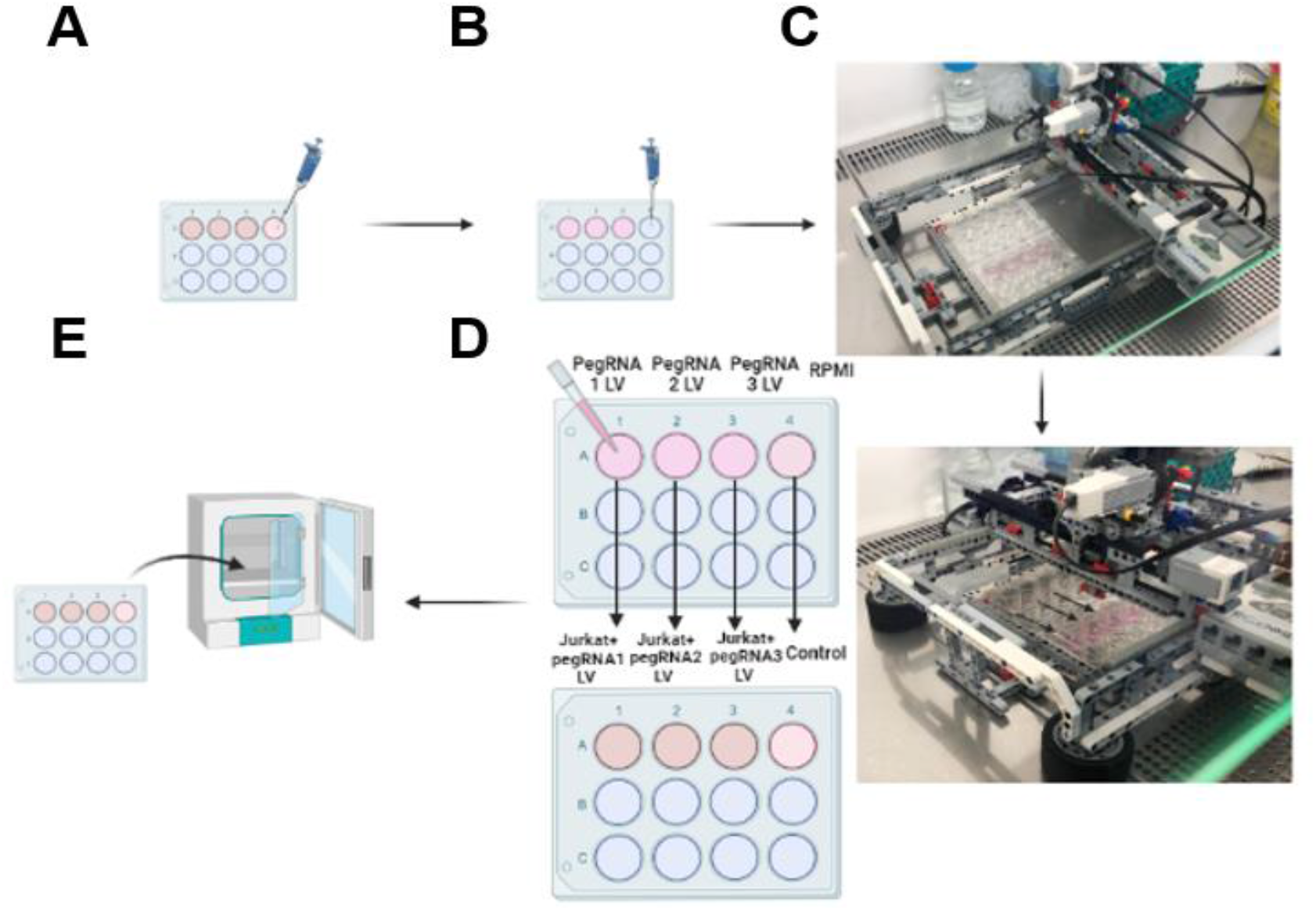
CRISPR Gene Modification. **A**. Seeding of Jurkat Cells into 12 well plate at 5×10^4^ cells/well. **B**. Virus Sample Preparation. gRNA 1,2,3 Lentivirus was placed in 12 well plate. **C**. Integration of Plates into the Autonomous Robot System, 12-well plates were placed in the frame of the Robot. **D**. Addition of gRNA Lentivirus to Cells. The wells containing gRNA Lentivirus 1,2,3 in the 12-well plate were drawn at 100° and added to wells 1, 2, and 3 separately. **E**. Incubation. After the robot performed the experiment, the plate was placed in the incubator.

With this experiment, it was observed that there was no virus expression in the control group, and gRNA1, gRNA2, and gRNA3 were added at 100 degrees angles, respectively, providing 35-40% GFP and guide RNA expression rate. Expression levels were determined by Flow Cytometry analysis on the 3rd and 7th days after the experiment was carried out (**Figure 20**). The results show that the CRISPR.BOT has the capability of editing cells using CRISPR encoding lentivirus.

**Figure 20.**
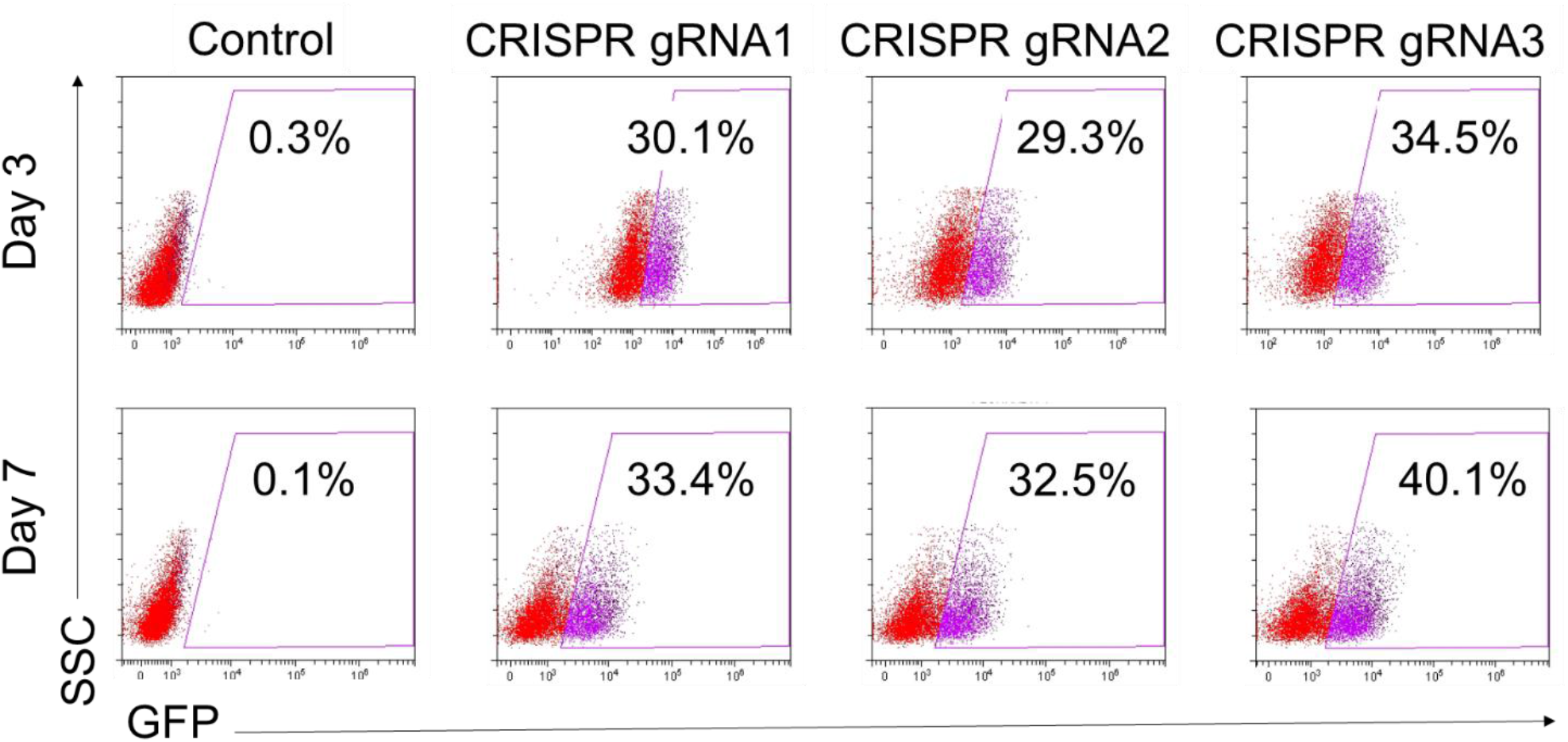
Flow Cytometry analysis of GFP protein expression in transgenic human cells encoding CRISPR.BOT modified CRISPR guide RNAs.

Optimization studies carried out for the pipette system in CRISPR.BOT V1 to effectively draw low microliter liquids have also been applied to the CRISPR.BOT V2 robotic system. With these studies, the sensitivity of the pipette system of CRISPR.BOT V2 and its ability to work with low microliters have been seen. Prior to the “Single-cell Sub-Cloning Procedures of Genetically Modified Cells” experiment, it was ensured that CRISPR.BOT V2 could perform Single-cell Sub-Cloning process with 96 well plate depending on the pipette precision and transfer appropriate microliters (**Figure 21**).

**Figure 21.**
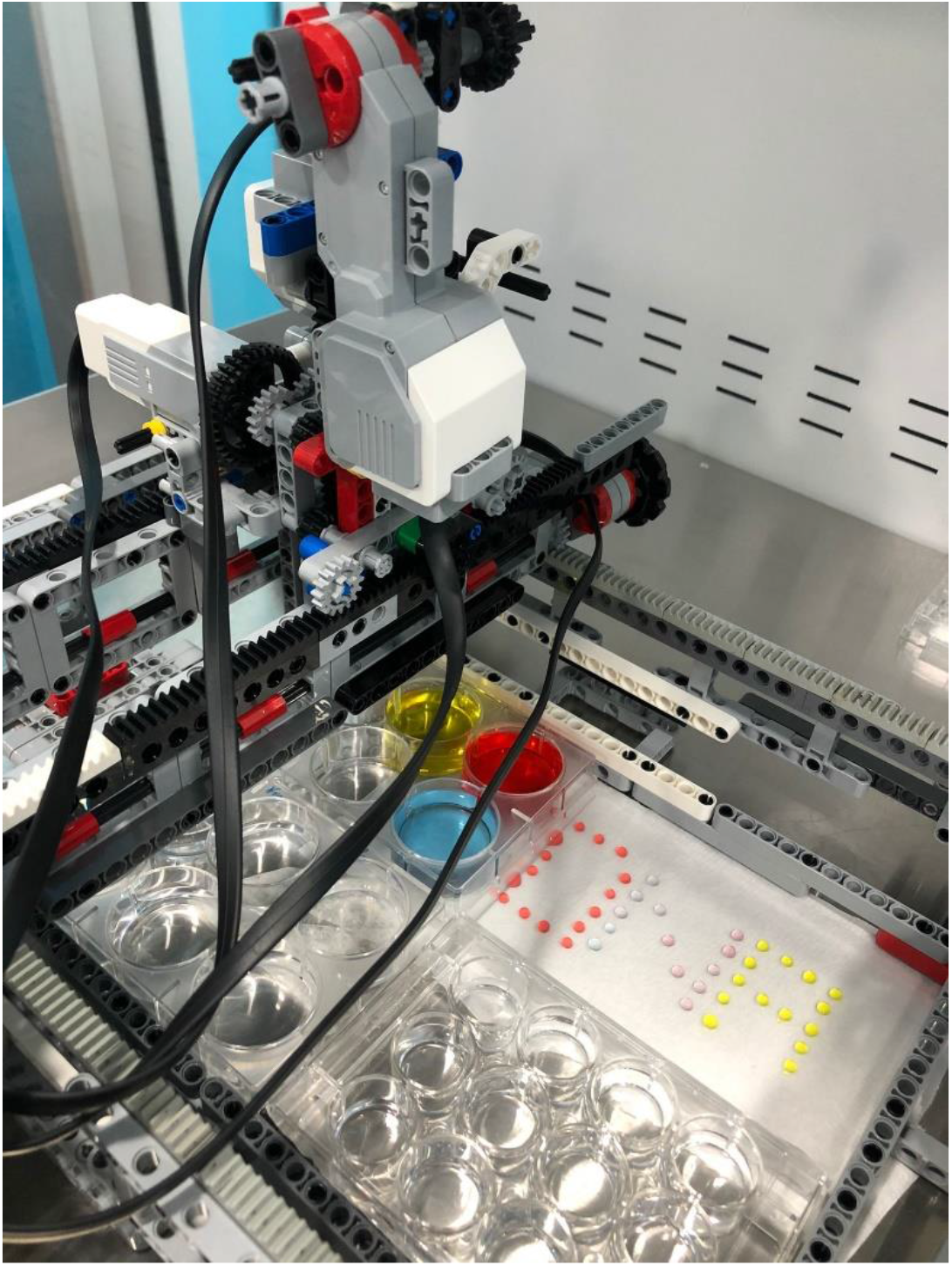
Precise microliter liquid intake performance trial studies with pipette system for CRISPR.BOT V2

#### Single-cell Sub-Cloning Procedures of Genetically Modified Cells

This experiment, it is aimed for the robot to drop the cells one by one without the need to use devices such as FACS. The main purpose of the evidential experiment for the future is to enable researchers to better understand the characteristics of a single cell population (subcloning) without the influence of other cells, with artificial intelligence-based autonomous systems. Jurkat cell encoding CRISPR guide RNAs were inoculated into 6 well plate with 5×10^4^ cells in each well. Prepared cell samples were placed on a 6-well plate and placed in the frame of the Robot. Then, the 96 well U Plate was placed on the frame of the Robot. Jurkat cells encoding CRISPR guide RNAs in 6 well plate at 100° were seeded into 72 wells of different 96 well U plate. After cultivation, the plates were transferred to an incubator at 37^0^C and 5% CO2 (**Figure 22**). For watching: https://drive.google.com/file/d/1B6elbjJYORL0gJE8GVw61Ux86fFs7IXx/view?usp=sharing

**Figure 22.**
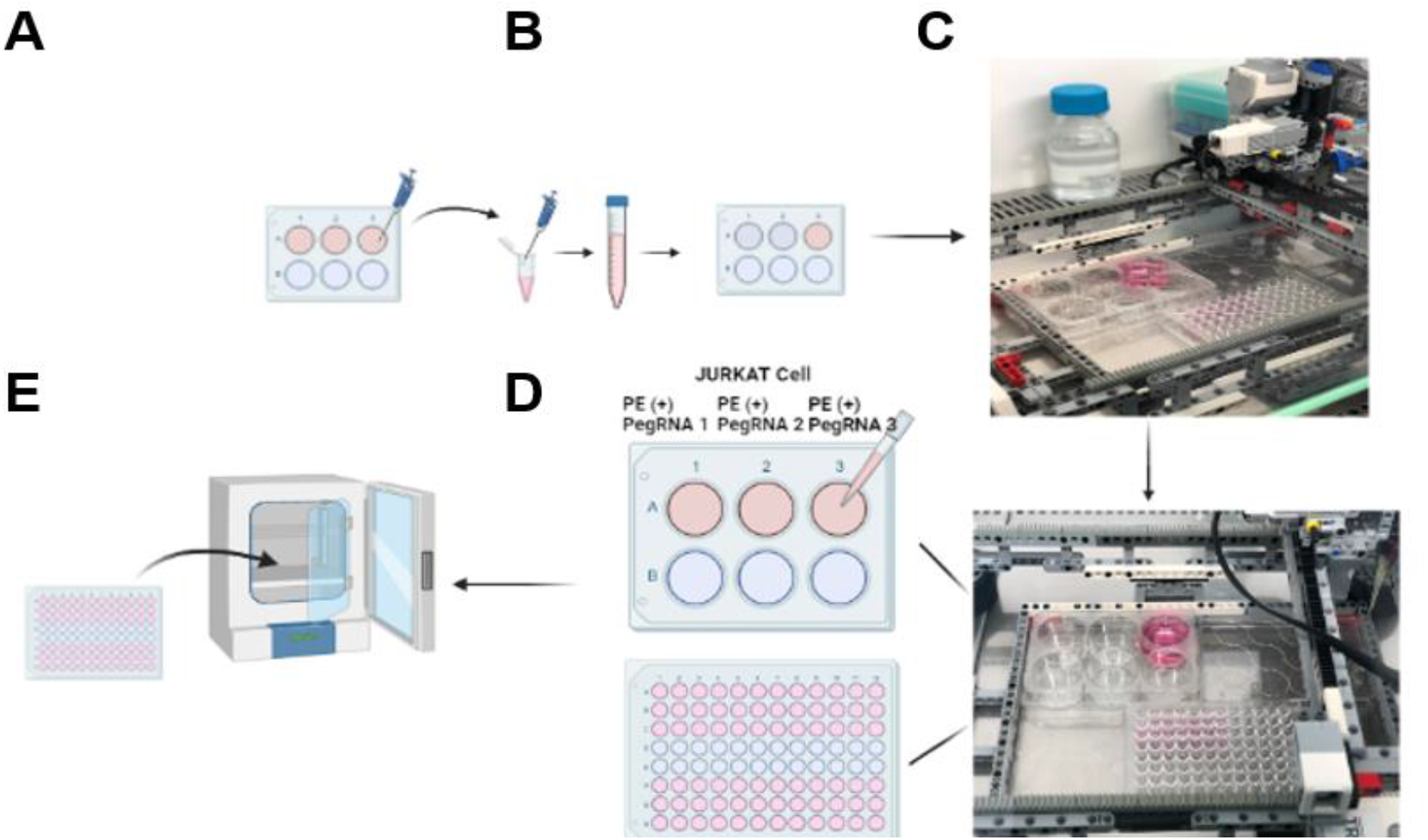
Single-cell Sub-Cloning Procedures of Genetically Modified Cells. **A**. Seeding of Cells. CRISPR + gRNA 1-2-3 Jurkat Cells were cultivated at 5×10^4^ cells/well in a 6-well **B**. Cell Dilution. 10µl of 3 different viruses in the wells of the 6-well plate were taken separately. 1 ml/500 cells were taken from the cell+RPMI mixture with a total volume of 1 ml and added onto 9 ml RMPI medium and placed in 6-well. **C**. Integration of Plates into the Autonomous Robot System. 6-well and 96-well plates were placed on the Robot’s frame. **D**. Sub-cloning of the cells to 96-well. CRISPR+ gRNA 1-2-3 Jurkat Cells in the 6-well plate were added to the wells in 96-well plates by pulling at 100°. **E**. Incubation of the 96-Well plate.

After 24 hours, plate scans were performed with an inverted microscope and cell photos were taken. After 7 days, plate scans were performed again with an inverted microscope and cell photographs were taken (**Figure 23**). These observations show that single or up to three-cell could be cultured per well as single cell sub cloning.

**Figure 23.**
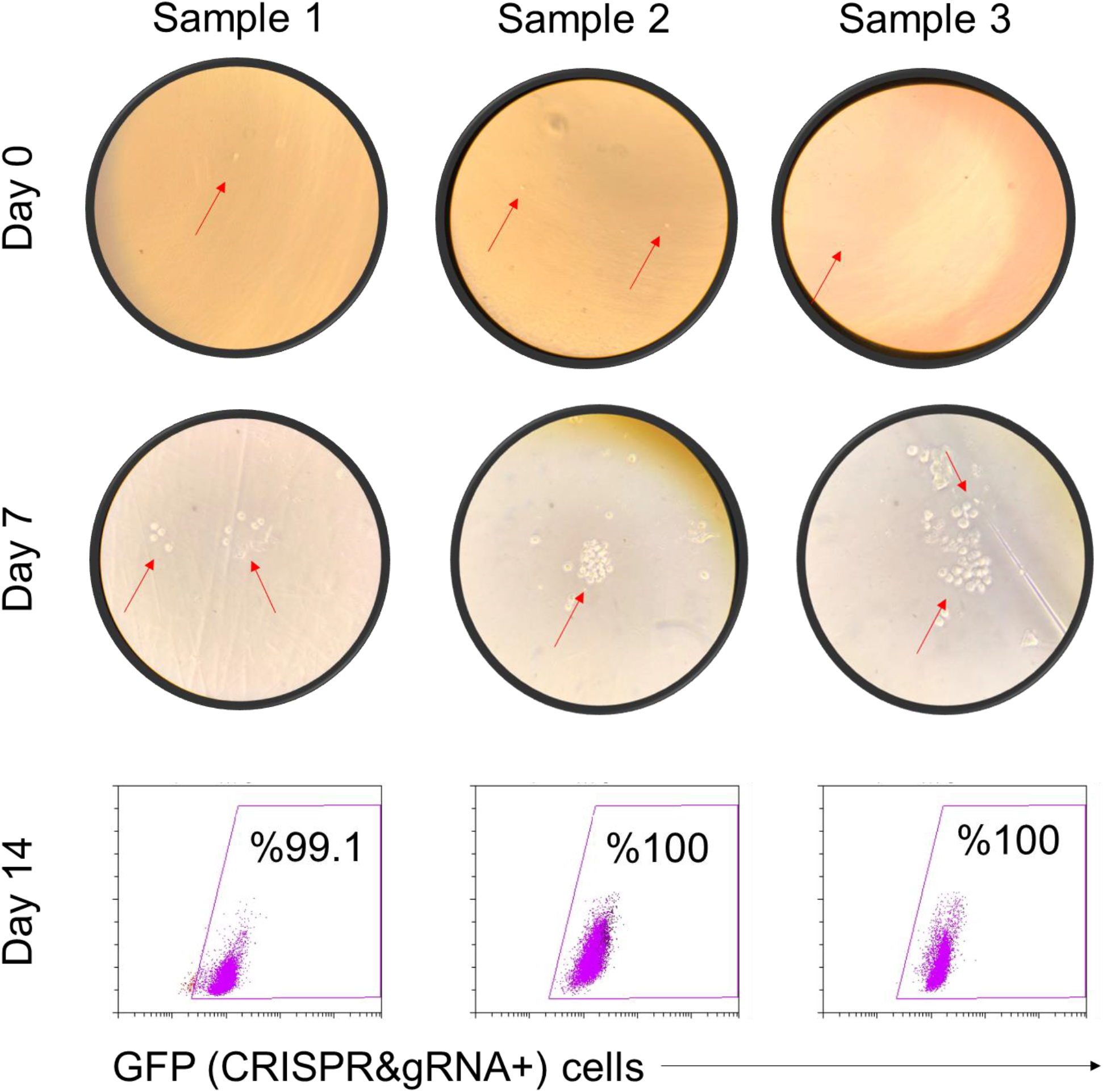
CRISPR.BOT-based subcloning of the transgenic cells.

Evaluation of the CRISPR.BOT-based subclones of transgenic cells, as seen in **Figure 24**, showed that the GFP expression was between 90-100% in the positive cell population and between 0-10% GFP expression in the negative cell population, the difference was clear, and a statistically significant value was found (**Figure 24**). These results showed that CRISPR.BOT accomplished single cell subcloning with a high efficiency and low cytotoxicity.

**Figure 24.**
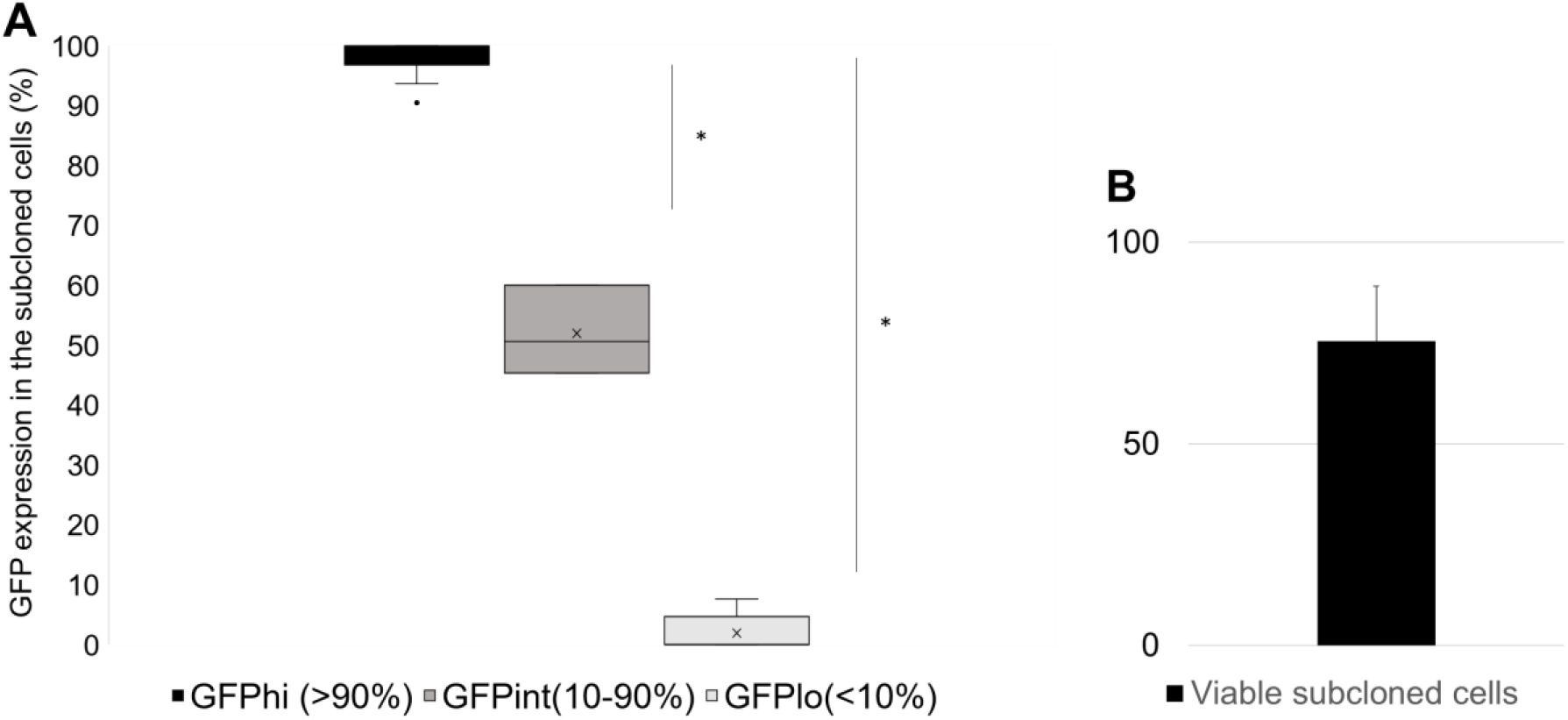
GFP expression and cell viability in subcloned cells. **A**. In the graph, the positivity rates of GFP expression of the sorted cells are GFPhi (>90%) and the negative rates are GFPlo (<10%). In the experiment performed with CROSPR.BOT, most of the samples were sorted and a statistically significant result was obtained. Intermediate level of GFP expression is seen in three of the samples, shown in the graph with GFPint (10-90%). **B**. The percentage of viability of subcloned cells is shown.

## Discussion

Liquid-processing robotic systems are also needed in life sciences laboratories to increase practicality and efficiency (Kong, 2012; Wohlsen, 2014). Today, systems that can handle automatic liquids and provide sample dispensing functions have become used in most life science laboratories (Chapman, 2003). However, human intervention does not disappear, it only reduces the risk of error. Robotic systems, on the other hand, can operate independently without the need for intervention at the start of the experiment (Tegally, 2020). It reduces the volume of the samples used in the experiments and increases the efficiency of the experiments both financially and in terms of workmanship (Kong, 2012). Studies of integrating automation systems with artificial intelligence (AI) provide the ability to optimize targeted therapeutic nanoparticles for unique cell types and patients (Egorov, 2021). Apart from the high costs of fully automatic systems, their maintenance is also quite expensive. There is a great cost loss because it usually requires system-specific protocols and laboratory equipment (Turbak et al., 2002). When we look at the market, there is no specialized robotic system that can perform reasonable molecular biology and genetic editing techniques that are affordable, sensitive, and at the same time meet the programmability requirements, and there is no robotic system that performs genetic engineering at the same time. The main purpose of the future-proof experiments carried out in the CRISPR.BOT project is machine learning, transgenic cell screening, analytical calculations, gene designs, genetic pathway selection, experimental design, cell cloning, and genetic pathway analysis through the development of remote and autonomous robotic systems in the field of molecular biology and genetics (**Figure 25**). It can be performed without human touch, with a very low margin of error. Gene transfer, genetic modification, and transgene cell cloning in bacteria and human cells were carried out autonomously for the first time with the algorithms we developed, with the CRISPR. In the future, these studies are thought to be useful for complex drug delivery experiments and many other applications in developing fields such as regenerative medicine and tissue engineering, in addition to biotechnological applications. Together with the CRISPR.BOT project, we have demonstrated that we can reduce human power and establish the first versions of an infrastructure in which molecular biology and genetic engineering can be done by robots in a closed system, without human involvement in pathogenic microorganisms (virus or bacteria, for example, SARS CoV-2 virus). With the results obtained from the experiments carried out, it is the prototype of a system in which viruses can be produced and used without human touch, and vaccine and drug studies can be carried out in studies to be carried out with viruses such as SARS-CoV2 for which inactive vaccine studies are dangerous. At the same time, with the development of this project, recombinant virus synthesis, characterization, testing, or predicting the experiment in vivo and in vitro, with the integration of artificial intelligence, which can be carried out autonomously with artificial intelligence and automated robotic experimental studies, plasmid isolation and offers significant potential competence to self-determine and perform many molecular biology experiments. In addition, a new direction will be given to the combination of artificial intelligence and robot, and a system will be created in which molecular biology and genetic methods can work 24 hours in a more automated, fast, and practical way.

**Figure 25.**
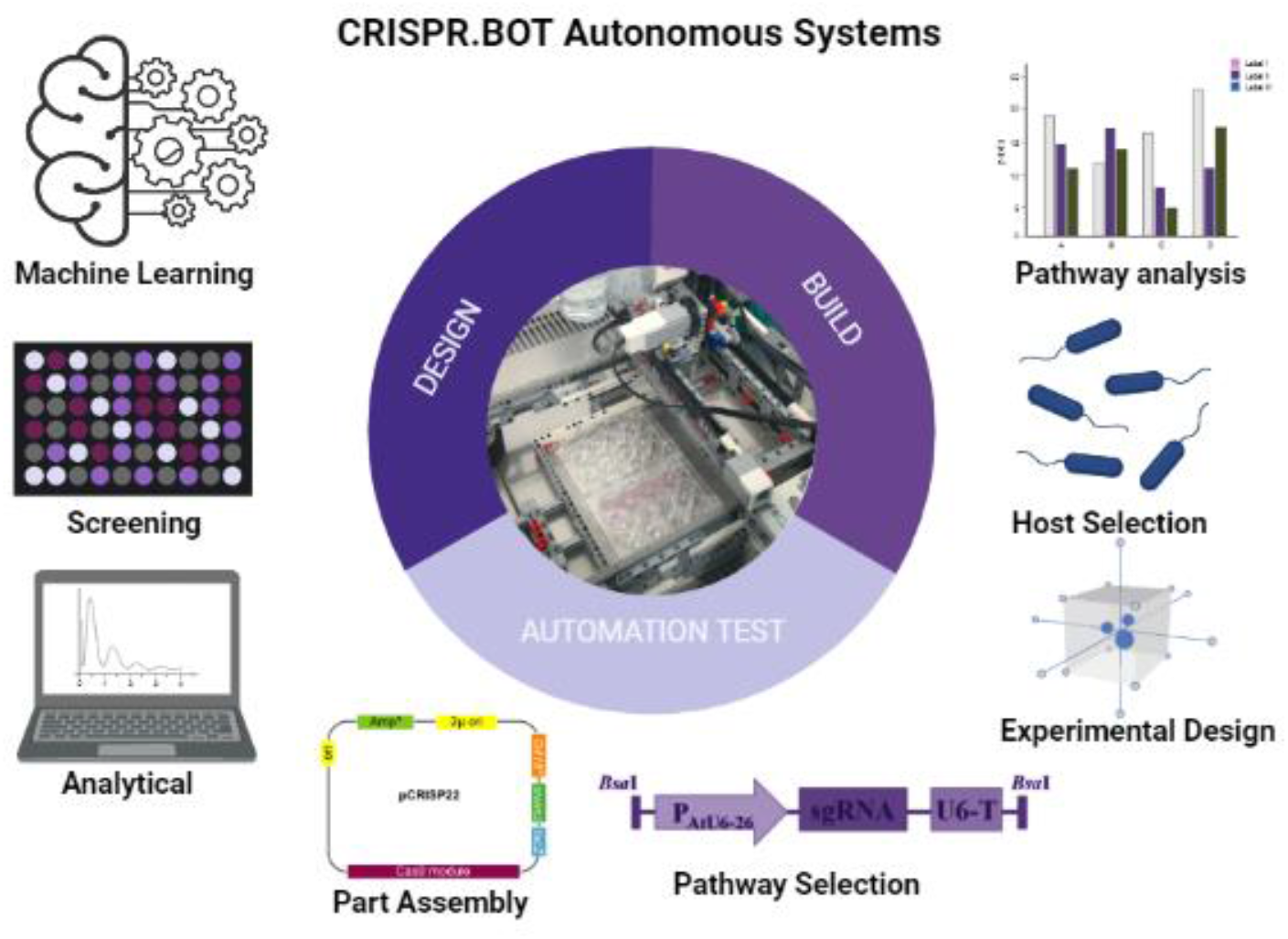
CRISPR.BOT autonomous systems. Experimental approaches that can be performed with CRISPR.BOT Autonomous Systems.

In addition to biotechnological applications in vaccine studies, transgenic cell generation such as recombinant viruses or dangerous viruses, genetic therapies, cancer studies, laboratory fields such as synthetic biology, microbiology or genetics, and in developing fields such as regenerative medicine and tissue engineering, complex drug delivery experiments and it is thought to be useful for many other applications (**Figure 26**).

**Figure 26.**
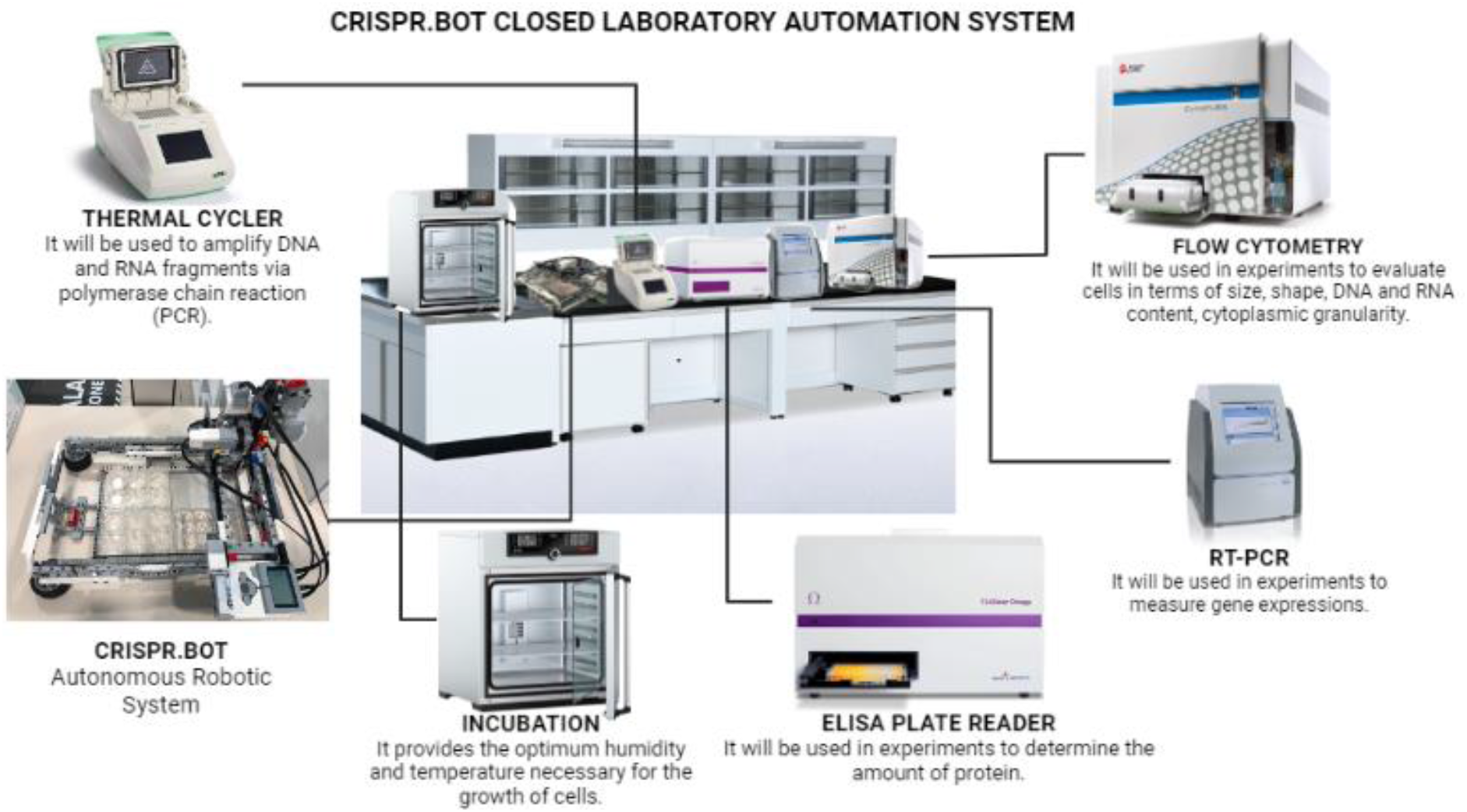
CRISPR.BOT closed laboratory automation system. Development of CRISPR.BOT Closed Laboratory Autonomous Systems and their integration with Laboratory analyzers (RT-PCR, ELISA Plate Reader, Flow Cytometry, Incubator, and Thermal Cycler).

The production of an end-to-end, high-efficiency laboratory automation system with ready-made workflow algorithms with CRISPR.BOT Robotic systems, of which we produced the prototypes, will enable us to establish a development company on molecular autonomous systems in the next stage. There are companies in the world (eg Miltenyl Prodigy, Lonza Cocoon, Molecular Devices) that manufacture devices based on existing automation and experimental customizations. It is anticipated that robotic systems that meet the application requirements of laboratories will increasingly continue in the near future. CRISPR.BOT Robotic systems will be a prominent development company in the future with the advantages of cost-effective, user-friendly, and remote control by researchers compared to their current competitors.

## Acknowledgment

We thank Prof. Dr. Kasif Nevzat Tarhan, Founder-Rector of Üsküdar University, Istanbul, Turkey for his vision and support genetic therapy field by founding TRGENMER Laboratory units in Üsküdar University in Istanbul. We thank LEGO System A/S, DK-7190 Billund, Denmark for providing several parts of LEGO Mindstorms to further develop CRISPR.BOT systems.

---

**Supplementary Figure 1.**
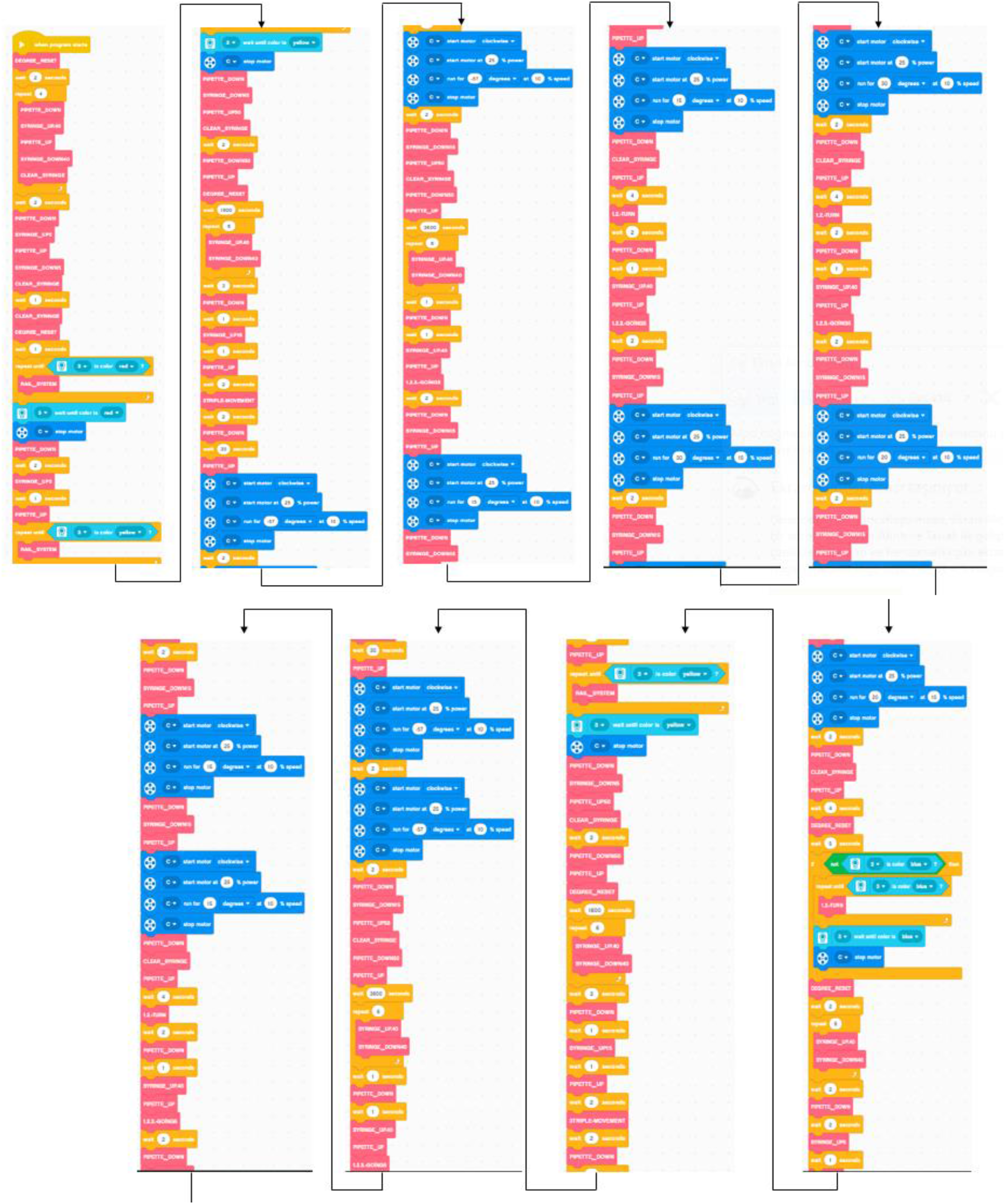
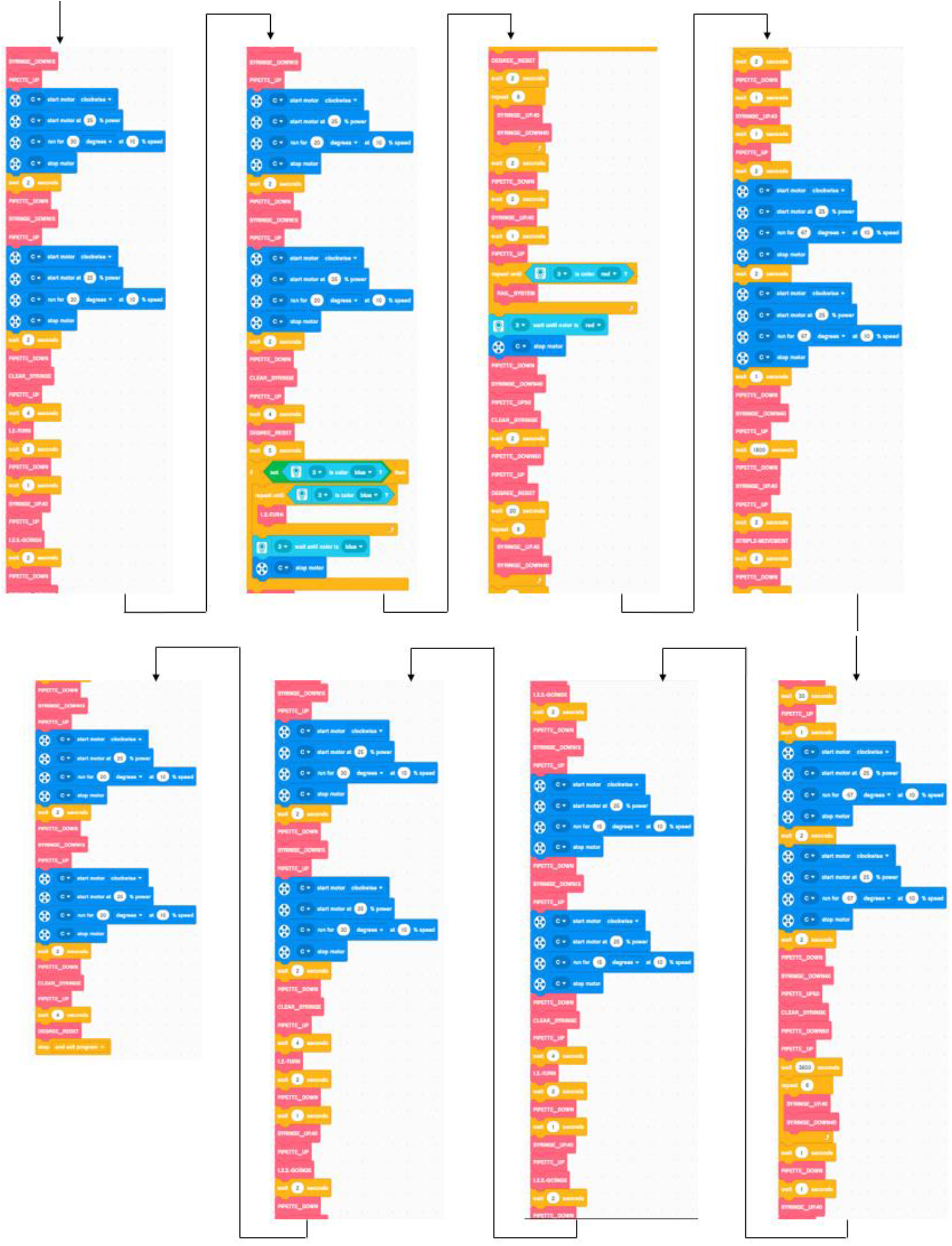
Bacterial transformation experiment programming. The degree reset block, it is aimed to reset the calibration of the robot before each experiment and to perform its movements more accurately. Before all trials, the robot was made to perform the newly added movements by resetting the degree. In order to measure the microliter of the liquid in the syringe, a syringe cleaning block is made and it is aimed to discharge the remaining liquids at the pipette tip and to get more accurate and sensitive results with this movement.

**Supplementary Figure 2.**
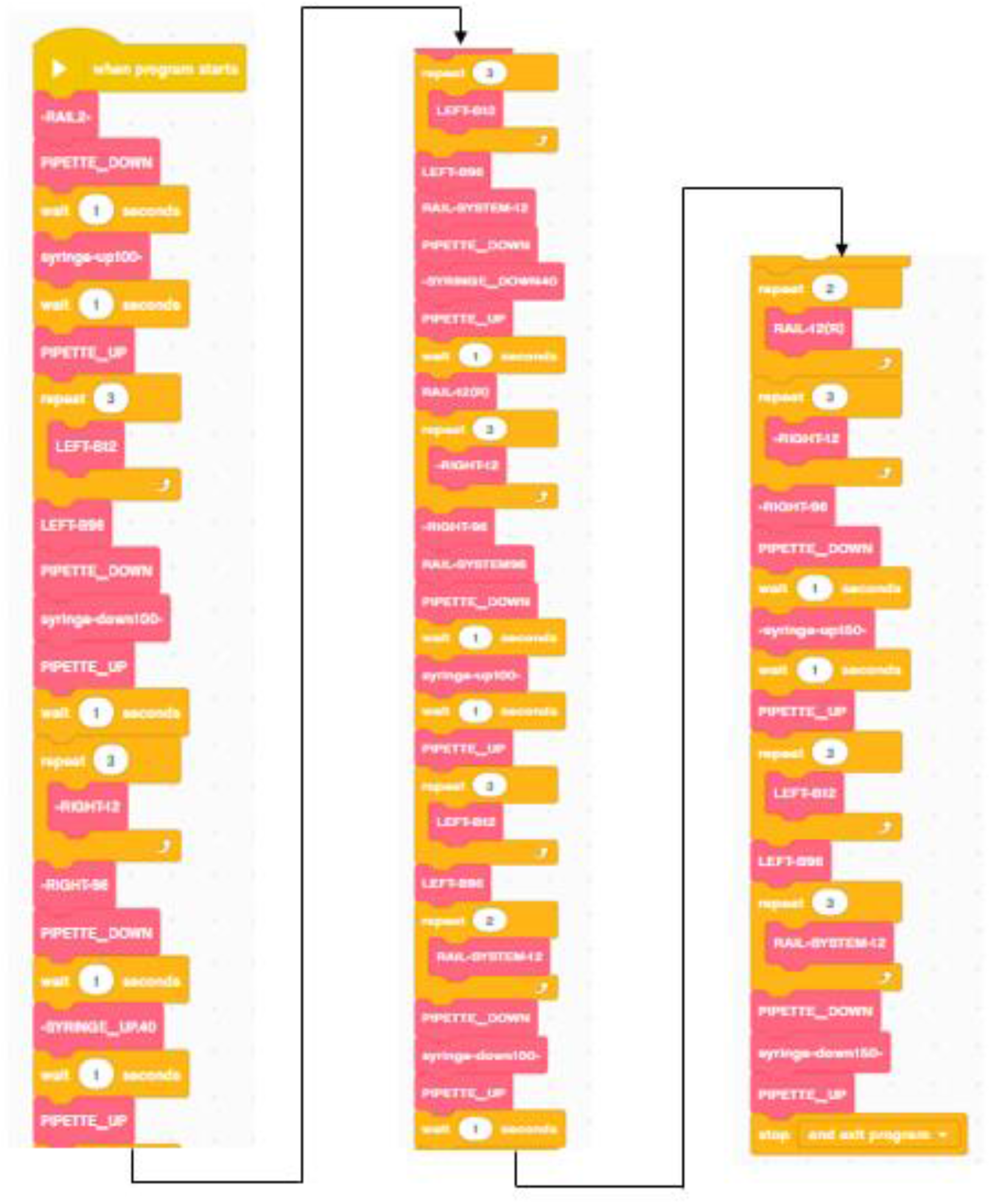
Programming that allows the robot to automatically genetically modify human cells. Rail and Rail-system blocks allow the robot’s pipette system to move from right to left or left to right on the frame. Left and Right blocks allow the pipette to pass between the cuvettes. For example, the Left-B12 block provides the movement of the pipette towards the brick on the 12 well plate. Grades are set specifically for the 12 well plate. Syringe-down and syringe-up blocks ensure that the syringe system can draw fluid. Pipette blocks, on the other hand, allow the syringe to enter and exit the wells, that is, up and down movements.

**Supplementary Figure 3.**
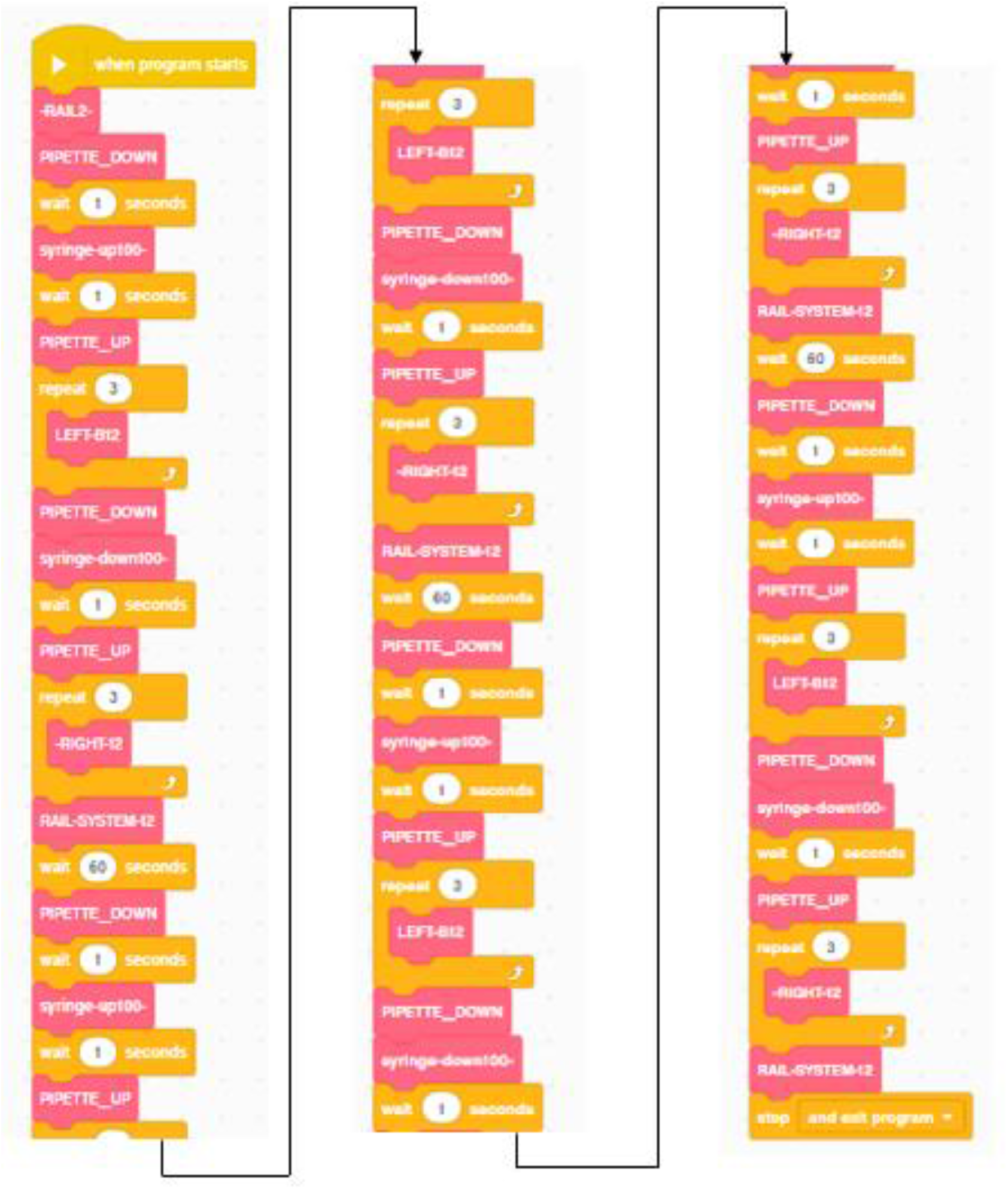
CRISPR Gene Modification experiment programming. With the experimental programming, it was aimed to transfer three different CRISPR guide RNAs of the robot to the cells. Movement commands are not defined as a block and are added to the programming by making calculations between plates for the desired movement. The places shown as 2 and 3 in the notes are the sequence of transfer of the guide RNA to the cells in the experimental phase.

**Supplementary Figure 4:**
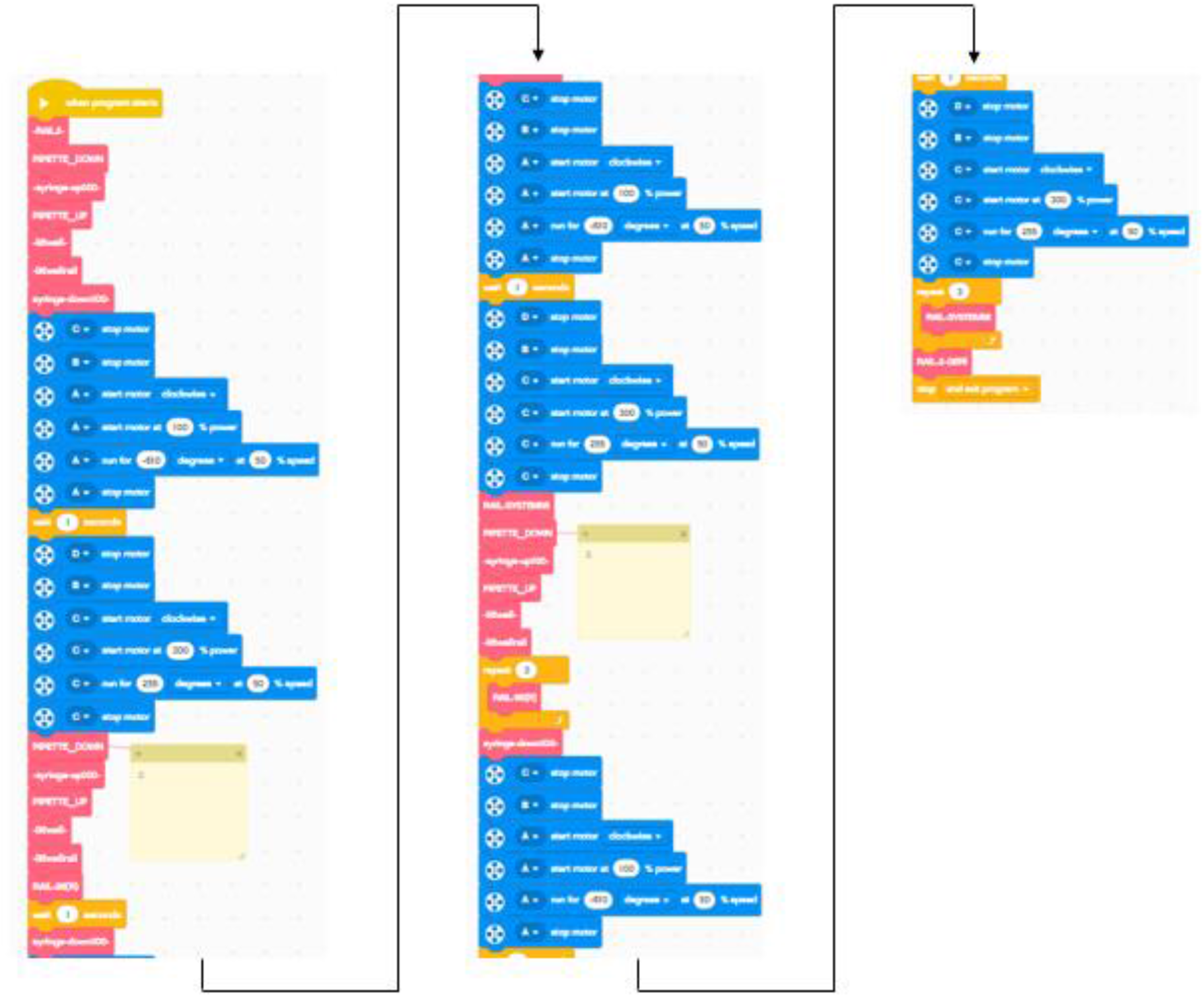
Cloning Procedures of Genetically Modified Cells Programing. With this experiment, it is aimed for the robot to drop the cells one by one without the need to use devices such as FACS or flow cytometry. Its programming is prepared to proceed in threes. It returns to the beginning every three wells and programming is started again.

